# Ribosome association primes the stringent factor Rel for recruitment of deacylated tRNA to ribosomal A-site

**DOI:** 10.1101/2020.01.17.910273

**Authors:** Hiraku Takada, Mohammad Roghanian, Julien Caballero-Montes, Katleen Van Nerom, Steffi Jimmy, Pavel Kudrin, Fabio Trebini, Rikinori Murayama, Genki Akanuma, Abel Garcia-Pino, Vasili Hauryliuk

## Abstract

In the Gram-positive Firmicute bacterium *Bacillus subtilis*, amino acid starvation induces synthesis of the alarmone (p)ppGpp by the multi-domain RelA/SpoT Homolog factor Rel. This bifunctional enzyme is capable of both synthesizing and hydrolysing (p)ppGpp. To detect amino acid deficiency, Rel monitors the aminoacylation status of the ribosomal A-site tRNA by directly inspecting the tRNA’s CCA end. Here we uncover the molecular mechanism of Rel-mediated stringent response. Off the ribosome, Rel assumes a ‘closed’ conformation which has predominantly (p)ppGpp hydrolysis activity. This state does not specifically inspect tRNA and the interaction is only moderately affected by tRNA aminoacylation. Once bound to the vacant ribosomal A-site, Rel assumes an ‘open’ conformation, which primes its TGS and Helical domains for specific recognition and recruitment of cognate deacylated tRNA to the ribosome. The tRNA locks Rel on the ribosome in a hyperactivated state that processively synthesises (p)ppGpp while the hydrolysis is suppressed. In stark contrast to non-specific tRNA interactions off the ribosome, tRNA-dependent Rel locking on the ribosome and activation of (p)ppGpp synthesis are highly specific and completely abrogated by tRNA aminoacylation. Binding pppGpp to a dedicated allosteric site located in the N-terminal catalytic domain region of the enzyme further enhances its synthetase activity.

## INTRODUCTION

Alarmone nucleotides guanosine pentaphosphate (pppGpp) and tetraphosphate (ppGpp), collectively referred to as (p)ppGpp, regulate metabolism, virulence, stress responses and antibiotic tolerance in the vast majority of bacterial species (for reviews see (1-4)). The cellular levels of (p)ppGpp are controlled by members of the RelA/SpoT Homolog (RSH) protein family (5). These enzymes both synthesise (p)ppGpp by transferring the pyrophosphate group of ATP onto the 3’ position of either GDP or GTP, and degrade the alarmone by hydrolysing the nucleotide back to GDP (or GTP), releasing inorganic pyrophosphate in the process.

RSH factors fall into two categories: ‘small’ single domain RSHs and ‘long’ multi-domain RSHs (5). In the majority of bacterial species, long RSHs are represented by a single ribosome-associated bifunctional enzyme called Rel that is capable of both synthesizing and degrading (p)ppGpp. In the lineage to Beta- and Gammaproteobacteria, duplication and functional divergence of the ancestral ribosome-associated Rel gave rise to a pair of specialized factors – the namesakes of the RSH protein family – RelA and SpoT (5,6). RelA is a synthesis-only enzyme incapable of hydrolysing (p)ppGpp (7), while SpoT and Rel are bifunctional (8,9). The (p)ppGpp synthesis activity of RelA and Rel is induced by so-called ‘starved’ ribosomal complexes (i.e. ribosomes accommodating deacylated tRNA in the A-site) (8,10). The association of SpoT with the ribosome is still a matter of debate (11,12). In the case of RelA, the pppGpp product is a potent inducer of the enzyme’s synthesis activity (13). The mechanistic basis of this regulatory mechanism is currently unexplored, and it is unknown whether bifunctional RSHs Rel and SpoT are similarly regulated by pppGpp.

Long RSHs are comprised of an enzymatic N-terminal multi-domain region (NTD) and a regulatory C-terminal multi-domain region (CTD) (**Figure 1A**). The NTD is subdivided into (p)ppGpp hydrolase (HD) and (p)ppGpp synthetase (SYNTH) catalytic domains, while the regulatory CTD is comprised of the TGS (ThrRS, GTPase and SpoT), Helical (equivalent to alpha-helical as per (14)), ZFD (Zinc Finger Domain; equivalent to CC, Conserved Cysteine as per (5) and RIS, Ribosome-InterSubunit as per (14)) and RRM (RNA recognition motif; equivalent to ACT, aspartokinase, chorismate mutase and TyrA, as per (5)). Experiments with Rel from *Streptococcus dysgalactiae* subsp. *equisimilis* (15,16), *Mycobacterium tuberculosis* (17,18), *Caulobacter crescentus* (19), *Staphylococcus aureus* (20) and *Thermus thermophilus* (21) have established that, first, the synthetic and hydrolytic activities of the NTD are mutually exclusive and, second, the CTD is essential for regulation by the ribosome.

**Figure 1.**
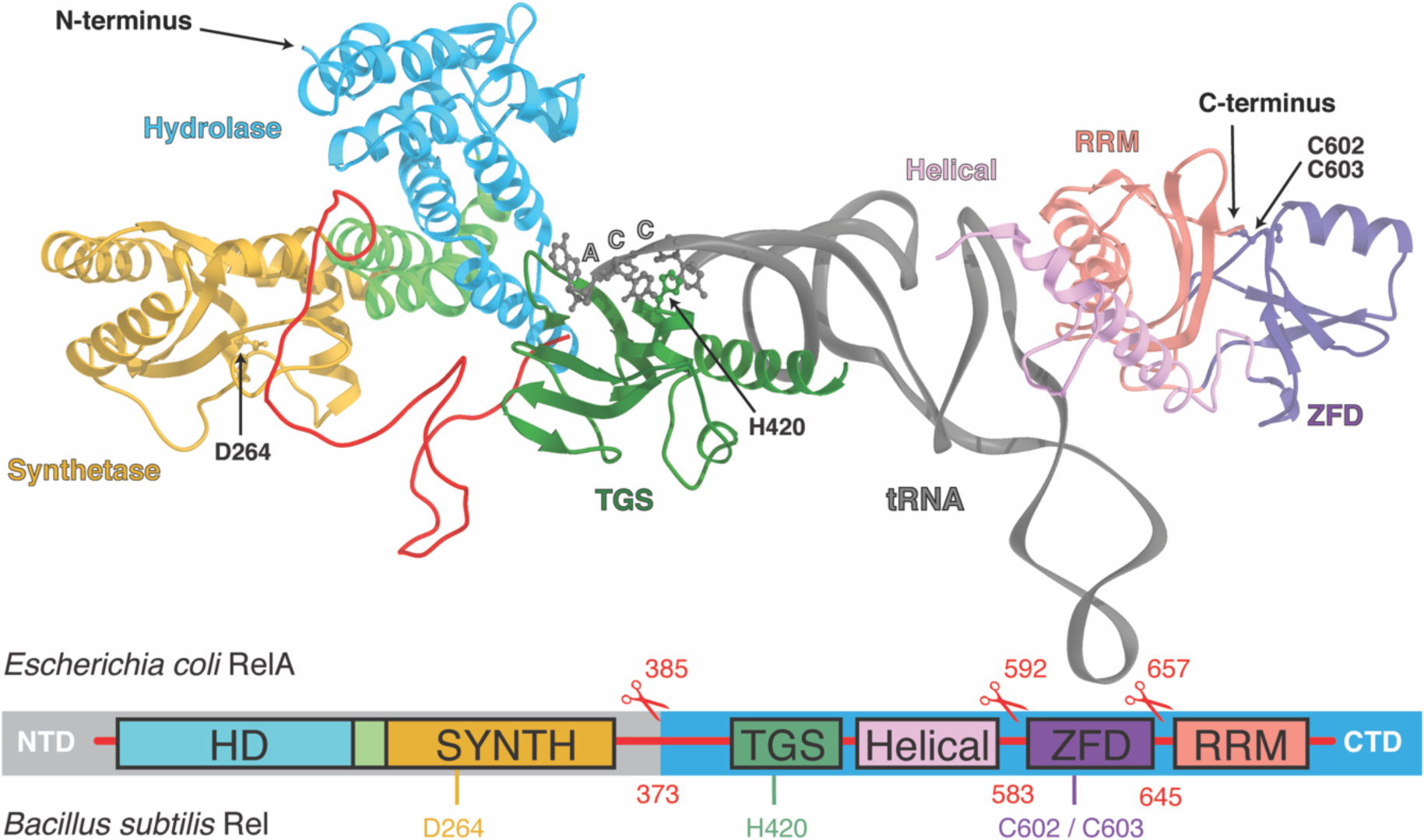
Molecular recognition of the A/R deacylated tRNA by the stringent factor. The domain composition as well as the truncated versions of the enzymes (*B. subtilis* Rel: ΔRRM 1-645, ΔZFD-RRM 1-583 and CTD 1-373; *E. coli* RelA: ΔRRM 1-657, ΔZFD-RRM 1-592 and CTD 1-385) used for functional studies are highlighted, as well as functionally important residues of *B. subtilis* Rel: D264, H420, C602-C603 and R619. The 3’ CCA end of A/R tRNA is shown as spheres. The 3D model was generated based on the structure by Loveland and colleagues (14), RDB accession number 5KPX.

Our understanding of the relationship between the enzymatic activity of RelA/Rel and the factors’ interactions with starved ribosomal complexes is largely based on experiments with *E. coli* RelA. A family of related ‘hopping’ models was put forward. According to the original ‘hopping’ model, each act of (p)ppGpp synthesis fuels the displacement of RelA from the ribosome, upon which the factor ‘hops’ to the next starved complex (22). Single molecule tracking experiments using RelA C-terminal fusions with photoswitchable fluorescent proteins led to the formulation of the ‘extended hopping’ (23) and the ‘short hopping time’ (24) models. The former model suggests that upon activation by starved ribosomes RelA processively synthesises (p)ppGpp off the ribosome and the latter states that upon activation, RelA spends prolonged periods of time catalytically active in association with starved ribosomal complexes. Finally, there are two ‘flavours’ of the ‘short hopping time’ model: RelA could first bind the vacant A-site and then recruit the deacylated tRNA (13,14) or form the RelA:tRNA complex off the ribosome and deliver deacylated tRNA to the A-site, similarly to how elongation factor EF-Tu delivers aminoacylated tRNA (25,26). The proposed delivery of deacylated tRNA to the ribosome by RelA/Rel is a controversial topic. While in the case of *M. tuberculosis*, Rel binding to tRNA is readily detectible (8,17), binding of tRNA to *E. coli* RelA is not detected by EMSA (13,26), unless the complex is crosslinked by formaldehyde (26). On the ribosome, the TGS and Helical domains of RelA form multiple contacts with deacylated A-site tRNA (14,27,28), but it is unclear if these domains are accessible when RelA/Rel is off the ribosome. Stable tRNA binding was reported in the case of a truncated version of *E. coli* RelA containing just the TGS and part of the Helical domain (26), suggesting that either the interaction interface is not accessible in the full-length protein or the interaction was not detected due to technical issues.

Studies of the physiological effects of (p)ppGpp in the Gram-positive Firmicute bacterium *Bacillus subtilis* have been fundamental to our understating of (p)ppGpp-mediated transcriptional regulation (29) and regulation of nucleotide metabolism (30). However, mechanistic understanding of *B. subtilis* Rel is lacking. Using our reconstituted biochemical system (31), we dissected the relationship between, on the one hand, Rel’s association with deacylated tRNA and the ribosome and, on the other hand, (p)ppGpp synthesis and hydrolysis by the enzyme, and probed the roles of individual domains. We establish that the specific recognition of deacylated tRNA takes place after Rel associates with the vacant ribosomal A-site, and the strength of the interaction with deacylated tRNA fine-tunes the stability of the interaction of Rel/RelA enzymes with starved ribosomes. In the case of *B. subtilis* Rel, the factor forms a stable complex with starved ribosomes and this complex is not actively dissociated by processive synthesis of ppGpp. We demonstrate that pppGpp enhances synthetase activity of both Rel and RelA by binding to a dedicated allosteric site located in the NTD region of the enzyme.

## MATERIALS AND METHODS

Detailed description of experimental procedures is provided in *Supplementary methods*. All bacterial strains and plasmids used in this study are listed in **Supplementary Tables 1** and **2**. All proteins were expressed, purified and characterised using SUMO-tagging strategy described earlier for *E. coli* RelA (32) and *B. subtilis* Rel (31); gel filtration profiles for mutant Rel variants are shown as (**Supplementary Figure S1**). Radiochemistry experiments were performed as described for *E. coli* RelA (13), with modifications. Sucrose gradient fractionation and Western blotting were performed as per (33), with minor modifications. Electrophoretic Mobility Shift Assay (EMSA) experiments were performed as per (34), with minor modifications. HPLC-based nucleotide quantification was performed as per Varik and colleagues (35). Wild type MG1655 and MG1655 *relA::HTF E. coli* (25) strains were grown in MOPS minimal media at 37 °C until OD_600_ 0.5 and challenged with 150 µg/mL of mupirocin (3X the MIC).

## RESULTS

### Off the ribosome the NTD region of B. subtilis *Rel non-specifically binds RNA*

When RelA is associated with the starved ribosomal complex, the TGS and Helical domains form specific contacts with the ribosome and tRNA (14,27,28). To probe the accessibility of Rel’s TGS and Helical domains for interaction with deacylated tRNA off the ribosome, we purified native, full-length RNA-free untagged *B. subtilis* Rel as well as a set of C-terminally truncated variants: lacking RRM (Rel^ΔRRM^, amino acids 1-645), lacking both RRM and ZFD (Rel^ΔZFD-RRM^, 1-583) and lacking all of the regulatory CTD domains (Rel^NTD^, 1-373). For the sake of simplicity, throughout the text *B. subtilis* Rel – the main focus of the current study – is referred to simply as ‘Rel’. In the case of Rel enzymes from other species, the organism is specified in subscript.

As a specificity control for tRNA binding studies we substituted a conserved histidine residue (H420E) in the TGS domain. The corresponding histidine residue in *E. coli* RelA (H432) forms a stacking interaction with the 3’ CCA end of the uncharged A-site tRNA (27). The H432E substitution abrogates RelA’s functionality by abolishing RelA activation by deacylated tRNA (25,31). We have shown a similar loss-of-function effect of the H420E substitution in *B. subtilis* Rel (31). Therefore, we concluded that the H420E substitution is an excellent tool to probe the specificity of Rel’s interaction with deacylated tRNA. We used the Electrophoretic Mobility Shift Assay (EMSA) to study complex formation between native deacylated *E. coli* tRNA^Val^ and *B. subtilis* Rel: full-length and C-terminally truncated, both wild type and H420E variants. While analogous EMSAs failed to detect a stable complex between *E. coli* RelA and deacylated tRNA (13,26), a *B. subtilis* Rel:tRNA complex is readily observable (**Figure 2A** and **Supplementary Figures S2**). In the absence of a non-specific RNA competitor, full-length Rel forms a complex with EC_50_ of 0.5 μM. Surprisingly, the H420E mutation decreases the affinity only twice (from EC_50_ 0.5 μM to 1.0 μM), suggesting a lack of specificity (**Figure 2A** and **Supplementary Figure S2AB**). The sequential deletion of RRM and ZFD domains increases the affinity to tRNA^Val^ (EC_50_ of 0.3 μM and 0.2 μM), and even upon deletion of the entire CTD, *B. subtilis* Rel^NTD^ affinity for tRNA^Val^ remains virtually unaffected (EC_50_ 0.7 μM) (**Figure 2B**, black filled trace, and **Supplementary Figure S2D**-**F**). Consistent with the isolated NTD binding tRNA^Val^, while the H420E substitution in the full-length and ΔZFD-RRM backgrounds does decrease the factor’s affinity to tRNA, it does not abrogate the interaction completely (**Figure 2B**, compare empty and filled traces). To further characterise the non-specific component of the Rel:tRNA interaction, we used single-stranded mRNA(MVF) as a competitor. In the presence of a 10-fold excess of mRNA over tRNA, the affinity for tRNA drops up to three-fold, demonstrating that Rel binds both RNA species with a similar affinity (**Figure 2B**, green traces, and **Supplementary Figure S3**). Using the set of truncated Rel variants described above, we the localized the source of this non-specific affinity for mRNA to Rel’s NTD region (**Figure 2C** and **Supplementary Figure S4**).

**Figure 2.**
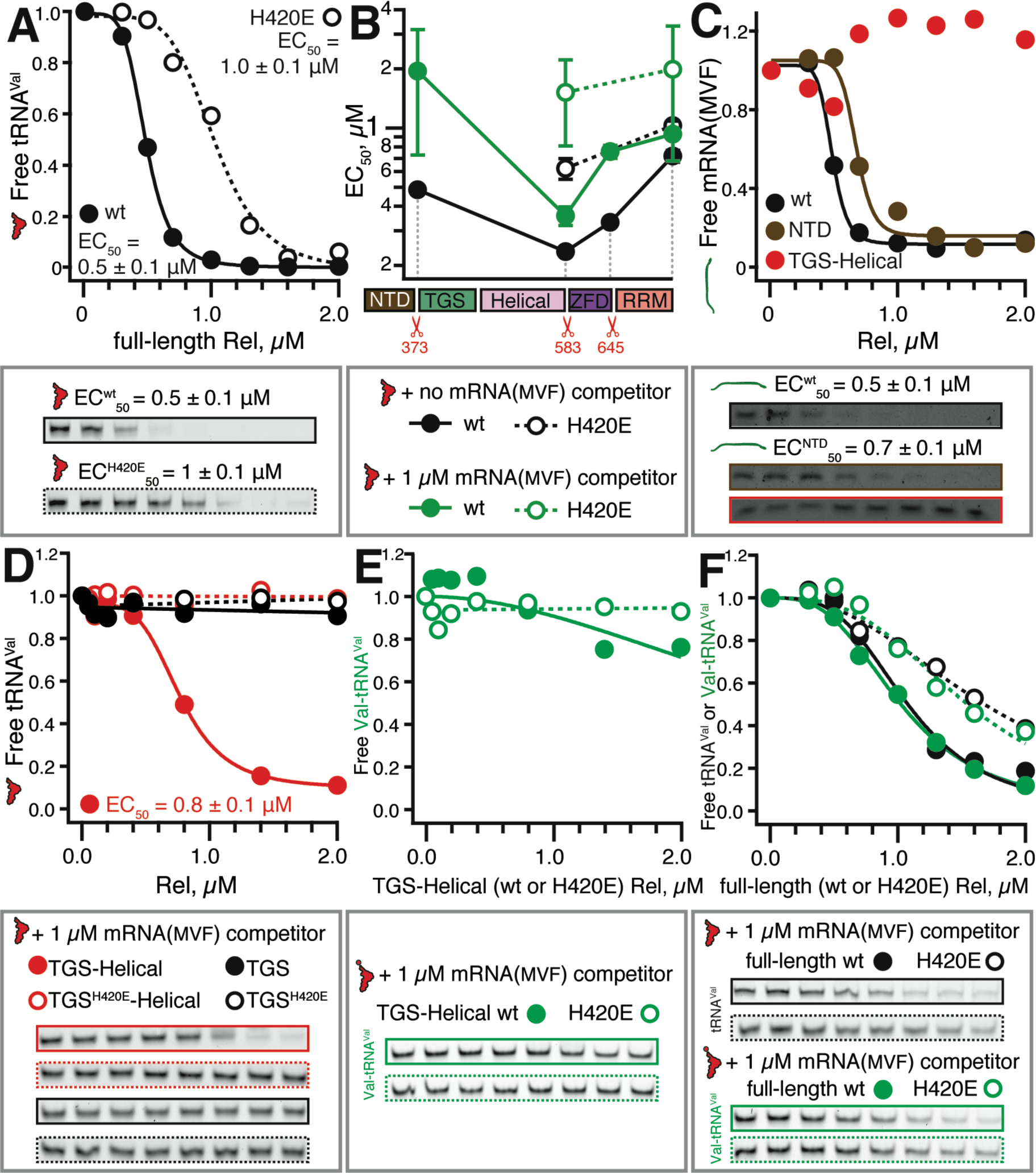
The isolated TGS-Helical region of *B. subtilis* Rel, but not the full-length protein, can specifically recognise the tRNA 3’ CCA end. Complex formation between increasing concentrations of *B. subtilis* Rel (wild type or mutants, H420E mutants shown as empty circles) and either 0.1 μM *E. coli* tRNA^Val^ or synthetic mRNA(MVF) was monitored by EMSA. Representative full EMSA gels as well as EC_50_ quantifications are provided as **Supplementary Figures S2**-**S5**. (**A**) Complex formation between *E. coli* tRNA^Val^ and either wild type or the H420E Rel mutant in the absence of a non-specific RNA competitor. (**B**) Complex formation between C-terminally truncated Rel variants and *E. coli* tRNA^Val^. (**C**) Complex formation between mRNA(MVF) model mRNA and full-length Rel, Rel NTD fragment or Rel TGS-Helical fragment. (**D**) Complex formation between *E. coli* tRNA^Val^ and either the Rel TGS domain or the Rel TGS-Helical fragment. Complex formation between acetylated and deacylated tRNA^Val^ and either isolated TGS-Helical fragment (**E**) or full-length (**F**) Rel. Analogous experiments using tRNA_i_^Met^ are presented on **Supplementary Figure S6**.

Taken together, these results suggest that in the absence of ribosomes, complex formation between Rel and tRNA is dominated by non-specific interactions mediated by the protein’s NTD region. This result is clearly at odds with the highly specific recognition of tRNA by the TGS domain in the cellular context as well as in in the presence of starved ribosomal complexes, both of which are abrogated by the H420E substitution (31). To deconvolute RNA binding by the NTD from that mediated by CTD, we next characterised the isolated *B. subtilis* Rel TGS-Helical region (amino acid positions 374-583) as well as the TGS domain alone (amino acid positions 374-469).

### The isolated TGS-Helical region of Rel specifically recognises the tRNA 3’ CCA end

While we do not detect formation of a stable complex in the case of TGS alone, the TGS-Helical fragment binds tRNA^Val^ with EC_50_ of 0.8 μM (**Figure 2D**). In stark contrast to the full-length protein, this interaction is completely abrogated upon introduction of the H420E mutation, suggesting that for these isolated regions (unlike the full-length protein), complex formation is driven by a specific recognition of the 3’ CCA tRNA end. Importantly, unlike the full-length enzyme and the NTD, the TGS-Helical fragment does not bind mRNA(MVF), further reinforcing the specificity of this interaction (**Supplementary Figure S5C**). Finally, aminoacylation of tRNA^Val^ completely abrogates the interaction with the isolated TGS-Helical domains (**Figure 2E**), while the affinity of the full-length Rel remains largely unaffected (**Figure 2F**). To rule out tRNA-specific effects, we performed an analogous set of binding assays with full-length Rel and the isolated TGS-Helical domains using both charged and deacylated *E. coli* initiator tRNA_i_^Met^ (**Supplementary Figure S6**). The results are in excellent agreement with our experiments with tRNA^Val^: while tRNA_i_^Met^ binding to full-length Rel is largely insensitive to the H420E mutation and tRNA aminoacylation, complex formation with isolated TGS-Helical domains is strictly dependent on deacylated tRNA_i_^Met^ and is abrogated by the H420E substitution.

Our results demonstrate that, unlike the full-length protein, the isolated Rel TGS-Helical region is highly specific in its recognition of the tRNA 3’ CCA end. To rationalise this result, we hypothesise that the association with the ribosome that drives the transition of Rel from the ‘closed’ conformation, unable to specifically sense and bind tRNA, to an ‘open’ conformation in which the TGS-Helical region is primed to specifically recognise the deacylated 3’ CCA end, thus driving Rel’s association with the starved ribosomal complex. Therefore, we next investigated the effects of deacylated tRNA on the interaction of Rel with the ribosome.

### Deacylated tRNA locks Rel on the ribosome in stationary phase B. subtilis

When *B. subtilis* cultures are challenged with the antibiotic mupirocin, deacylated tRNA^Ile^ accumulates in the cell and the bulk of Rel is efficiently recruited to the ribosome (**Figure 3A**). The effect can be readily reconstituted biochemically using purified Rel and *B. subtilis* 70S initiation complexes (**Supplementary Figure S7**). The A-site tRNA-dependent Rel recruitment to starved complexes is highly specific and is efficiently abrogated by tRNA aminoacylation.

**Figure 3.**
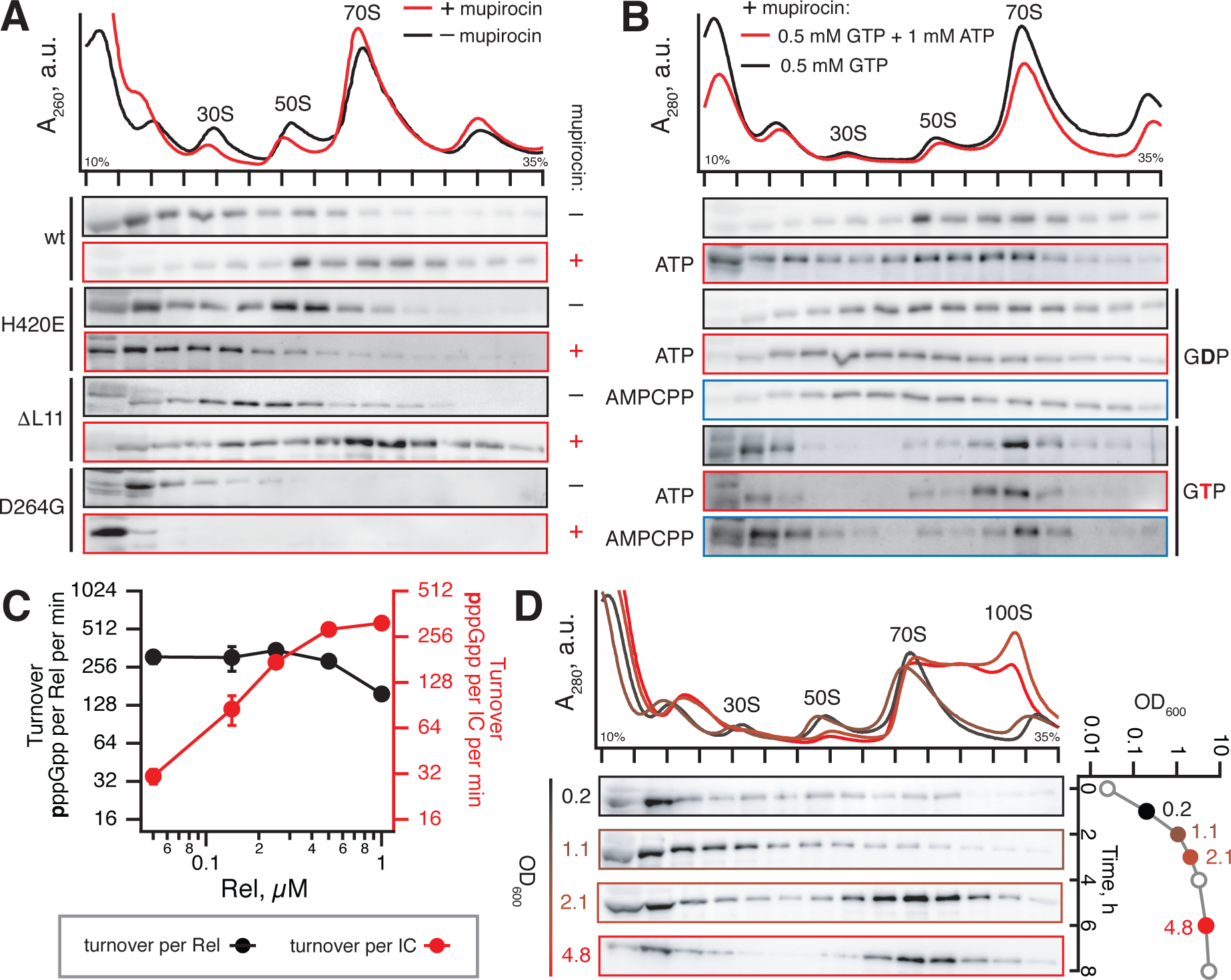
Rel is stably recruited to the starved ribosome upon amino acid starvation to drive processive (p)ppGpp production. (**A**) Polysome profile and immunoblot analyses probing Rel’s interaction with the ribosomes with or without the induction of acute isoleucine starvation by antibiotic mupirocin. Wild type Rel was expressed in either wild type 168 *B. subtilis* or in ΔL11 strain (VHB47). Rel H420E variant defective in CCA recognition was expressed from the native chromosomal locus in wild type *B. subtilis* (VHB68). Synthesis-inactive D264G was ectopically expressed in Δ*rel B. subtilis* under the control of P_*hy-spnak*_ promotor (VHB156), and expression was induced by 1 mM IPTG. (**B**) Effects of nucleotide substrates on Rel’s association with starved ribosomes generated by *B. subtilis* by treatment with mupirocin. (**C**) Synthase activity of Rel activated by starved ribosomal complexes (0.5 μM 70S IC(MVF) and 2 μM tRNA^Val^) as a function increasing concentrations of Rel. Error bars represent standard deviations of the turnover estimates by linear regression and each experiment was performed at least three times. (**D**) Ribosomal association of Rel as a function of *B. subtilis* growth phase. All experiments were performed with *B. subtilis* grown in liquid LB media at 37 °C.

We tested the association of Rel with ribosomes in conditions that naturally activate the stringent response, as opposed to acute non-physiological isoleucine starvation caused by mupirocin (**Figure 3D**). As bacteria enter the stationary phase and nutrients become limiting, the ribosome-associated RSHs RelA/Rel are activated and the intracellular concentration of (p)ppGpp increases (35). We characterised Rel association with ribosomes in *B. subtilis* throughout the growth curve, collecting the samples at OD_600_ of 0.2, 1.1, 2.1 and 4.8. In the last two samples, the bulk of Rel is recruited to ribosomes and the 100S ribosome dimer peak is prominent. The latter is a sign of high (p)ppGpp levels: expression of the ribosome dimerization factor HPF is under positive stringent control (36) resulting in formation of 100S ribosome dimers upon entry to the stationary phase (37).

### *E. coli* RelA does not stably associate with starved ribosomal complexes due to low affinity to deacylated tRNA

Despite multiple contacts between tRNA and RelA on starved ribosomal complexes (14,27,28), our earlier EMSAs using full-length *E. coli* RelA failed to detect a stable complex with deacylated tRNA off the ribosome (13). At the same time, Kushwaha and colleagues have recently reported nM-range tRNA affinity of the *E. coli* RelA fragment containing the TGS domain and part of the Helical domain (26). Therefore, we tested tRNA binding to a set of C-terminal truncations of *E. coli* RelA (**Supplementary Figure S8A**), as well as the isolated TGS-Helical fragment (**Supplementary Figure S8B**). However, we did not detect stable complex formation in either of the cases. While the reason for this discrepancy is unclear, our EMSAs strongly suggest that *E. coli* RelA is a considerably weaker tRNA binder than *B. subtilis* Rel.

Since deacylated tRNA is the driving force promoting association of the stringent factor with the ribosome, this implies that *E. coli* RelA association with starved ribosomes is, correspondingly, less stable as well. We assessed this interaction using sucrose gradient centrifugation of lysates, with and without mupirocin pre-treatment. To detect RelA by Western blotting we used an *E. coli* strain encoding RelA C-terminally tagged with His_6_-TEV-FLAG (HTF) on the chromosome. This tagged RelA displays wild type-like functionality in live cells (**Supplementary Figure S9**) (25). While the bulk of RelA is shifted to denser fractions upon amino acid starvation, the protein does not form a stable complex (**Figure 5A**), demonstrating that *E. coli* RelA is a significantly weaker binder of starved complexes compared to *B. subtilis* Rel.

### Recruitment of Rel to ribosomes by deacylated tRNA is not sufficient to induce synthesis activity in the absence of L11

To dissect the relationship between Rel recruitment to the ribosome and (p)ppGpp synthesis, we used three mutant strains with compromised (p)ppGpp synthesis activity (**Figure 3A**). First, we tested the effect of mupirocin on Rel carrying the H420E substitution that abrogated ribosomal recruitment in the reconstituted system. In contrast to wild type Rel, we detect no ribosomal recruitment of H420E Rel. Similarly, no tRNA-dependent ribosomal recruitment of H420E Rel was observed with reconstituted 70S initiation complexes (**Supplementary Figure S7**). Next, we tested Rel with a D264G substitution in the SYNTH domain, which abrogates (p)ppGpp synthesis by the enzyme (38). Surprisingly, D264G Rel does not bind ribosomes regardless of the presence or absence of mupirocin (**Figure 3A**). This could be explained by the allosteric coupling between both catalytic domains: by disrupting GDP/GTP substrate binding, the D264G substitution could be driving the protein into the SYNTH^OFF^ HD^ON^ state thus destabilising its binding to the ribosome. Indeed, our ^3^H-pppGpp hydrolysis assays support this hypothesis (see below and **Figure 4E**).

**Figure 4.**
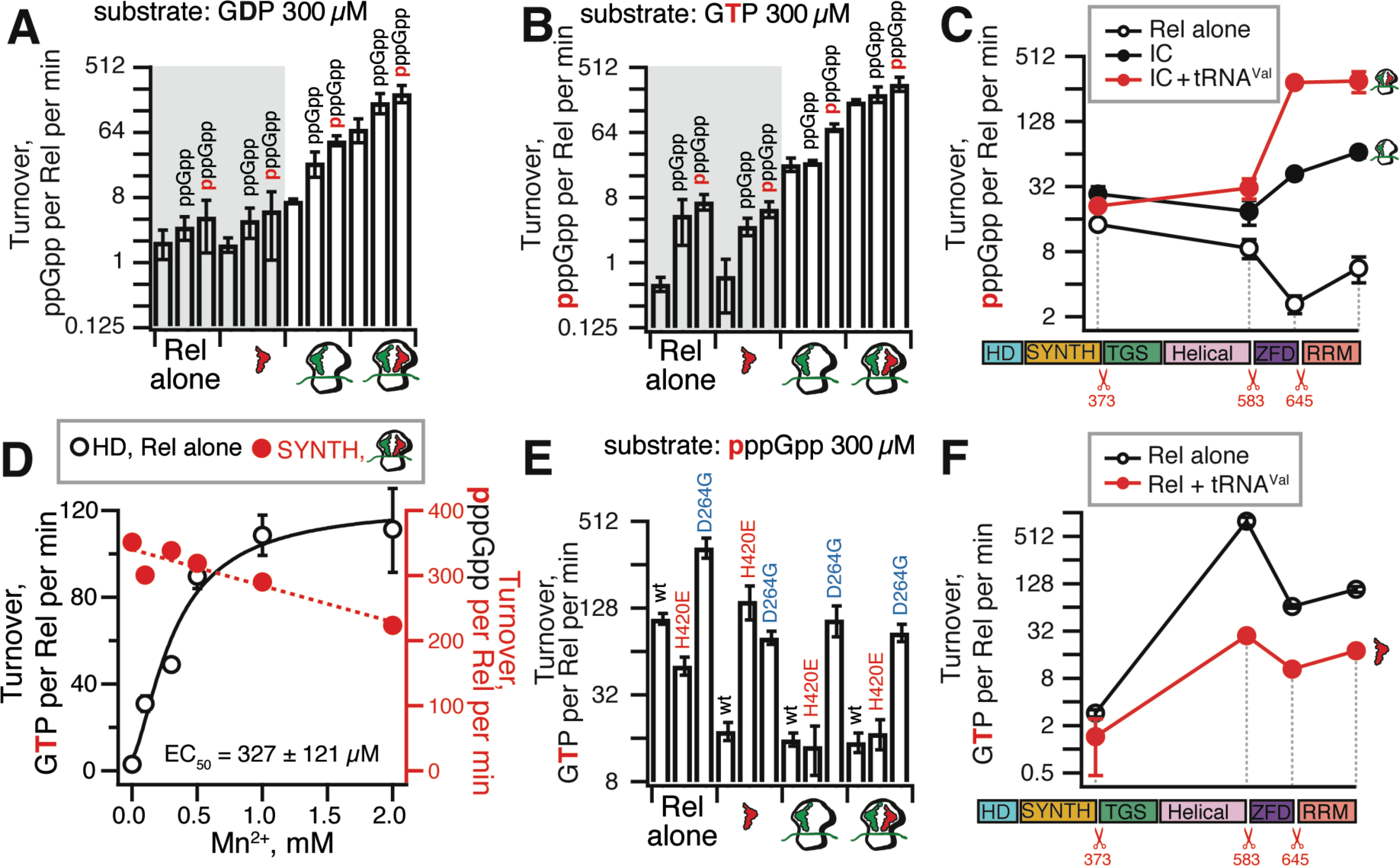
Role of individual domains in the regulation of Rel’s (p)ppGpp synthesis and hydrolysis activities by tRNA, ribosomes, starved ribosomal complexes and pppGpp. Synthase activity of 140 nM wild-type and C-terminally truncated Rel assayed in the presence of 1 mM ATP (**A**-**C**) and 0.3 mM of either ^3^H-labelled GDP (**A**) or GTP (**B** and **C**). Hydrolase activity of 140 nM wt and C-terminally truncated Rel variants assayed in the presence of 0.3 mM of ^3^H-labelled pppGpp (**D**-**F**). As indicated on the figure, the reaction mixtures were supplemented with combinations of 0.5 μM IC (MV) and tRNA^Val^ (2 μM: A-site) as well as 100 μM ppGpp (**A, B** and **D**) or pppGpp (**A**-**C**). All experiments were performed in HEPES:Polymix buffer, pH 7.5 at 37 °C in the presence of 5 mM Mg^2+^ (**A**-**C**) and additionally in the presence of either increasing concentrations of Mn^2+^ (**D**) or 1 mM Mn^2+^ (**E** and **F**). Error bars represent standard deviations of the turnover estimates by linear regression and each experiment was performed at least three times.

Finally, we tested a *B. subtilis* strain lacking ribosomal protein L11. This protein is crucial for activation of *E. coli* RelA by starved ribosomal complexes, both in live bacteria and in the reconstituted biochemical system (7,22,39), as well as for the functionality of *C. crescentus* Rel (40). The lack of L11 does not perturb Rel’s recruitment to the ribosome upon mupirocin treatment, demonstrating that association of Rel with starved complexes *per se* is not sufficient for activation of its synthetic activity.

### Processive (p)ppGpp synthesis by ribosome-associated Rel does not induce ‘hopping’

Next, we reassessed the ‘hopping’ models by testing the effects of nucleotide substrates for (p)ppGpp synthesis on the interaction of wild type Rel with starved ribosomes in cell lysates (**Figure 3B**). We supplemented both the lysates and the corresponding sucrose gradients with guanosine substrates (0.5 mM GTP or GDP), both added alone or together with either ATP or its non-hydrolysable analogue, α,β-methyleneadenosine 5’-triphosphate (AMPCPP, 1 mM). This set of conditions was used to discriminate between the effects of substrate binding *per se* and active catalysis, since the latter was suggested to actively fuel the dissociation of *E. coli* stringent factor RelA from the ribosome (22). In the presence of the GDP substrate, the Western blot signal of Rel is spread out in the ribosomal fractions, with further addition of ATP or AMPCPP having only a minor effect (if any). In the case of GTP, Rel is detected both in the light fractions (free protein) and in the 70S ribosomal peak. Just as in the case of GDP, further addition of either ATP or AMPCPP does not have a significant effect. Taken together, these results suggest that the catalytic activity – or the lack of it (i.e. upon omission of ATP or its substitution for AMPCPP) – does not play a significant role in Rel association with starved ribosomes. There is, however, a clear difference between the effect of GDP and GTP on Rel association with ribosomes. This effect could arise from the GTP-dependent action of other A-site binding factors present in lysates, e.g. translational GTPases or different conformations induced by each nucleotide.

To further discriminate between ‘hopping’ and processive synthesis on the ribosome, we titrated wild type Rel in our reconstituted biochemical system. When the reaction turnover is calculated per starved ribosomal complex (**Figure 3C**, red trace), the enzymatic activity reaches a plateau when the concentration of Rel is equal to that of the ribosomes (0.5 μM). At higher Rel concentrations the efficiency of (p)ppGpp production in the reconstituted system does not increase – and when turnover is calculated per Rel molecule, it decreases (**Figure 3C**, black trace). This behaviour is consistent with Rel processively synthesising (p)ppGpp while associated with starved complexes rather than the enzyme spending prolonged periods off the ribosome in a catalytically active state upon departure from the ribosome. In the latter case one could expect that, acting catalytically, one starved ribosomal complex would fully activate several Rel molecules; this is not the case.

### The hydrolysis substrate pppGpp does not promote Rel dissociation from starved ribosomal complexes

Given the well-documented antagonistic allosteric coupling between the SYNTH and HD catalytic domains of Rel enzymes (16,21), we tested the effect of the hydrolysis substrates ppGpp and pppGpp on Rel’s interaction with the ribosomes. However, due to the significant volume of sucrose gradients (12 mL) it is not feasible to perform the experiment in a way that makes the substrate available for the enzyme as it migrates though the gradient. Therefore, we resorted to supplementing only the lysate with 1 mM pppGpp. While under these conditions we see no effect on Rel’s association with the ribosome (**Supplementary Figure S10**), there are at least two possible explanations for this result. First, pppGpp could be diluted in the test tube and thus lose its effect. Second, the association of Rel with the starved complexes could inhibit pppGpp binding to the HD domain, thus counteracting the effect of this substrate. Our enzymatic experiments directly support the latter explanation (**Figure 4E**, see below).

### The synthetase activity of Rel is activated by pppGpp binding to an allosteric site in the NTD region

Next, we characterised the effects of substrates (GDP or GTP) as well as regulators (ribosomal complexes, tRNA, and pppGpp) on the synthesis activity of Rel. Since Mn^2+^ is universally essential for hydrolysis activity of long RSHs such as *M. tuberculosis* Rel (8), *S. equisimilis* Rel (15) and *E. coli* SpoT (41), we characterised the synthesis activity of the full-length Rel in the absence of divalent manganese ions in order to avoid possible underestimation of the synthesis efficiency due to concomitant (p)ppGpp hydrolysis.

While the ribosome stimulates Rel’s synthesis activity 5-to 10-fold, and the ultimate activator – the starved complex – has a significantly stronger effect, approximately 50-fold (**Figure 4AB**). In good agreement with our earlier results with *E. coli* RelA (13), deacylated tRNA by itself has no significant effect. As with Rel enzymes from *S. equisimilis* (15) and *M. tuberculosis* (18,42), *B. subtilis* Rel is moderately more efficient in converting GTP to pppGpp than converting GDP to ppGpp. Conversely, *E. coli* RelA prefers the GDP substrate (13,42). The preference to GTP likely plays a regulatory role since it is GTP consumption, rather than direct (p)ppGpp binding to RNA polymerase that effectuates the transcriptional regulation upon the stringent response in *B. subtilis* (29,43). Finally, similarly to *E. coli* RelA (13), pppGpp stimulates *B. subtilis* Rel synthetic activity and ppGpp has a significantly weaker effect.

Stimulation of the synthesis activity of *B. subtilis* Rel by pppGpp implies complex formation between the alarmone and the protein. We used isothermal titration calorimetry (ITC) to study the binding of pppGpp to the NTD-only variant of Rel which is dramatically more soluble than the full-length protein and, just like the full-length protein, is activated by pppGpp (**Supplementary Figure S11A**). The Rel NTD region binds pppGpp with an affinity of 10.6 ± 0.9 μM (**Table 1** and **Supplementary Figure S11B**). Importantly, the interaction has a stoichiometry close to unity, i.e. one Rel molecule binds one pppGpp molecule. To test the possibility of the heat signal being reflective of the alarmone binding to one of the two active sites, we titrated pppGpp into Rel^NTD^ in the presence of saturating concentrations of GDP and a non-hydrolysable ATP analogue AMPCPP (to prevent (p)ppGpp biding in the catalytic pocket of the SYNTH domain), or GDP and AMPCPP combined with EDTA (to remove the Mn^2+^ ion from the catalytic site of the HD domain and prevent the potential biding of (p)ppGpp by this site). In both cases, both the affinity (*K*_*D*_) or the stoichiometry (*n*) of pppGpp binding remained largely unperturbed. Taken together, our results suggest the existence of a dedicated pppGpp-binding allosteric site located in the NTD region of Rel.

**Table 1.**
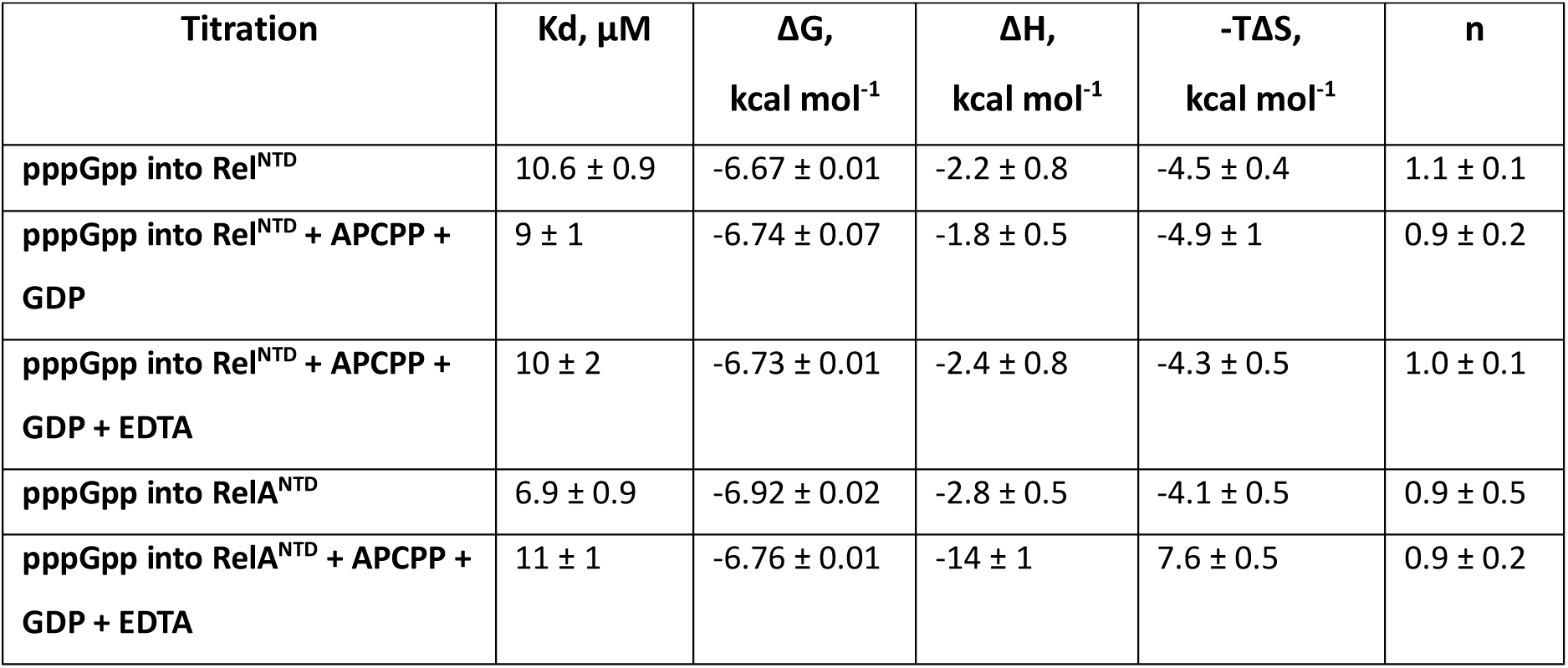
Thermodynamic parameters of pppGpp binding to NTD domain region fragments of *B. subtilis* Rel and *E. coli* RelA as determined by ITC. The binding affinities were determined from fitting a single interaction model to the experimental ITC data. Data represent mean values ± standard deviations.

### The pppGpp-binding site is conserved between Rel and RelA

To test if, similarly to *B. subtilis* Rel^NTD^, the allosteric pppGpp binding site of *E. coli* RelA is also located in the NTD region, we performed a set of enzymatic assays with C-terminally truncated *E. coli* RelA variants in the presence and absence of 100 µM pppGpp (**Figure 5BC**). Deletion of the RelA RRM domain compromises activation by the A-site tRNA of the starved complex (31). A likely explanation is that the low affinity of RelA to starved complexes renders it more sensitive to destabilisation of the complex by deletion of the RRM domain. Activation by the ribosome itself is, however, refractory to progressive deletion of the regulatory CTD. Importantly, when CTD-truncated RelA is assayed in the absence of ribosomal complexes, we do not detect the increase in its synthetic activity that would be expected in the auto-inhibition model (15,44,45).

**Figure 5.**
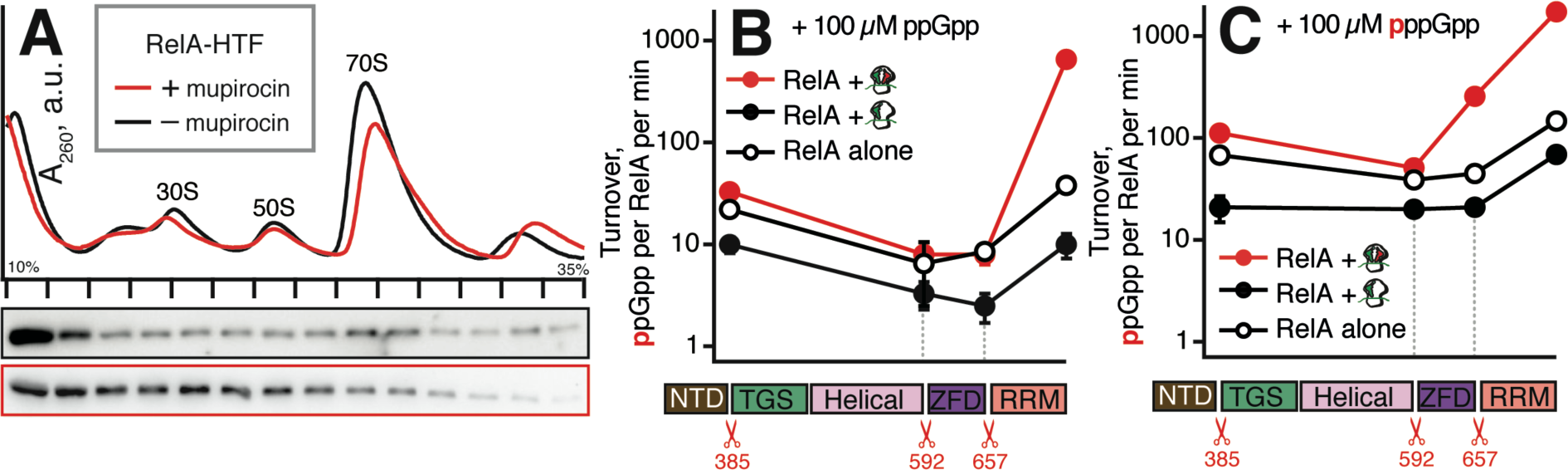
*E. coli* RelA NTD region contains the allosteric regulatory site that binds pppGpp. (**A**) MG1655 *E. coli* (*relA*::*HTF*) expressing was expressing C-teminally HTF-tagged RelA was grown in LB liquid medium at 37 °C. To induce amino acid starvation bacterial cultures exponentially growing at OD_600_ 0.2 were treated for 20 min with mupirocin added to final concentration of 70 µM that completely abolishes bacterial growth. (**B** and **C**) Synthesis activity of wild type *E. coli* RelA as well as RelA^ΔRRM^ (1-657), RelA^ΔZFD^ (1-592), and RelA^NTD^ (1-385) mutants by assayed in absence of ribosomes, in presence of 70S initiation complexes and 70S initiation complexes supplemented with cognate deacylated tRNA^Val^. Experiments were performed in the presence of either 100 µM ppGpp (**B**) or pppGpp (**C**). Error bars represent standard deviations of the turnover estimates by linear regression and each experiment was performed at least three times.

Importantly, all of the truncated variants of *E. coli* RelA are consistently activated by pppGpp and, similarly to Rel, this holds even for the NTD-only version. The strength of the effect varies from 2 to 32-fold depending on the construct and whether the protein is tested alone or in the presence of 70S initiation or starved complexes. Direct measurements by ITC confirmed that pppGpp directly binds to RelA^NTD^ with an affinity of 6.9 ± 0.9 μM, and is insensitive to the addition of GDP, AMPCPP and EDTA (**Table 1** and **Supplementary Figure S11EF**). Taken together, our results demonstrate that as with *B. subtilis* Rel, the allosteric pppGpp-binding allosteric regulatory site of *E. coli* RelA is located in the NTD region of the enzyme.

### Starved ribosomal complexes actively induce Rel’s synthesis activity rather than merely release the auto-inhibition by CTD

Next, we used enzymatic assays in a reconstituted biochemical system to probe the roles of the individual domains in sensing the ribosome and A-site deacylated tRNA. Deletion of the RRM domain had little effect on activation by starved complexes and pppGpp (**Figure 4C** and **Supplementary Figure S12**). The activation by initiation complexes was compromised, but not abrogated, and the synthetic activity of the Rel^ΔRRM^ alone was weaker than that of the wild type. These results are seemingly at odds with mis-regulation and over-production of (p)ppGpp by both Rel^ΔRRM^ (19) and RelA^ΔRRM^ (32) in live cells, which could be attributed to the loss of autoinhibition in the mutant protein (15,18,44). This interpretation is not supported by our biochemical results, and follow-up experiments established that the toxicity of both *B. subtilis* Rel^ΔRRM^ and *E. coli* RelA^ΔRRM^ is strictly dependent on the interaction with starved ribosomes (31).

Progressive removal of RRM and ZFD domains abrogates activation by A-site tRNA, although activation by the initiation complex itself remains detectable. In good agreement with earlier results obtained with *M. tuberculosis* Rel (17,18), *B. subtilis* Rel NTD has a synthesis activity that is only about two-fold higher than the full-length protein. This demonstrates that release of the autoinhibition of the NTD by the CTD does not account for the full activity observed in the presence of starved ribosomes.

### The ribosome and tRNA inhibit (p)ppGpp hydrolysis by Rel

The synthetic and hydrolytic activities of Rel’s NTD are mutually exclusive (15,16,21). This motivated us to directly test the effects of the ligands that induce the synthetic activity (ribosomes, tRNA) on Rel’s hydrolysis activity. The hydrolase activity of the enzyme tested alone is strictly Mn^2+^-dependent and peaks at 1 mM of Mn^2+^(**Figure 4D**). We tested the effects of tRNA, ribosomes and starved complexes (**Figure 4E**). In good agreement with earlier results with *M. tuberculosis* Rel (8), tRNA inhibits the hydrolysis activity of *B. subtilis* Rel, and the inhibition is abrogated when the interaction with the tRNA CCA end is disrupted by the H420E substitution. Both initiation and starved complexes inhibit hydrolysis, and the H420E substitution does not overcome this effect. Therefore, we conclude that it is the ribosome itself that inhibits hydrolysis. Synthesis-inactive D264E Rel has elevated hydrolysis activity, consistent with the antagonistic allosteric coupling between SYNTH and HD domains (16,21). This activity is insensitive to inhibition by ribosomes, in good agreement with the ribosomal recruitment being compromised by the mutation (**Figure 3A**).

Next, we tested the HD activity of our C-terminally truncated Rel variants, both in the presence and absence of tRNA^Val^ (**Figure 4F**). Progressive deletion of both RRM and ZFD leads to induction of the hydrolysis activity (**Figure 4F**), while the synthesis activity is compromised (**Figure 4C**). Reduction of Rel to an NTD fragment lacking the regulatory CTD near-completely abrogates the hydrolysis activity, in good agreement with our microbiological (31) and ITC experiments (**Figure 5**), as well as biochemical studies of *S. equisimilis* (15) and *C. crescentus* (19) Rel enzymes. The inhibitory effect of tRNA^Val^ is lost only when Rel is reduced to its hydrolytically near-inactive NTD.

Taken together, our biochemical results demonstrate that the CTD-mediated transduction of the regulatory stimuli that induce (p)ppGpp synthesis inhibits the hydrolysis activity – and *vice versa*. This regulatory strategy in the full-length protein utilises the antagonistic allosteric coupling within the NTD enzymatic core of the protein and efficiently prevents futile enzymatic activity.

## DISCUSSION

Almost half a century after Haseltine and Block demonstrated that *E. coli* RelA is activated by cognate deacylated tRNA in the ribosomal A-site (10), the exact molecular mechanism by which long ribosome-associated RSHs sense amino acid starvation is still unresolved. The long list of outstanding questions incudes: does Rel bind the tRNA off the ribosome – and recruits it to the empty A-site – or does it bind the ribosome first? Are synthesis and hydrolysis compatible with stable recruitment of Rel to starved ribosomal complexes: can ribosome-associated Rel hydrolyse (p)ppGpp? can it processively synthesyse the alarmone or does Rel ‘hop’? What is the exact role of the individual domains in, on one hand, recognition of tRNA and the ribosome and, on the other hand, intra-molecular regulation of the two opposing enzymatic activities? Is the synthesis activity of Rel stimulated by pppGpp analogously to that of RelA, and if yes, where is the allosteric pppGpp binding site located in long ribosome-depended RSH enzymes?

Here we tackle these questions through extensive comparative functional characterisation of *B. subtilis* Rel and *E. coli* RelA. We propose a model that summarises our results (**Figure 6**). Off the ribosome, Rel adopts a ‘closed’ conformation in which tRNA-binding TGS and Helical domains are sequestered. In this state the protein has net hydrolase activity, i.e. HD^ON^ SYNTH^OFF^. Amino acid starvation depletes the pool of ternary complexes formed by aminoacylated tRNA associated with elongation factor EF-Tu in a GTP-bound state. Rel samples the vacant ribosomal A-site and assumes an ‘open’ conformation, thus presenting its tRNA-binding TGS and Helical domains. In this conformation, (p)ppGpp hydrolysis is supressed, and (p)ppGpp synthesis is moderately activated (HD^OFF^ SYNTH^ON^). Subsequent binding of the cognate tRNA stabilises Rel on the ribosome and serves as the ultimate inducer of (p)ppGpp synthesis (HD^OFF^ SYNTH^TURBO^). The product of the reaction, pppGpp, binds to a dedicated allosteric site enzyme located in the NTD region of both Rel and RelA, and serves as an inducer of the (p)ppGpp synthesis activity. Individual acts of catalysis are not incompatible with prolonged association with the ribosome, and multiple turnovers are performed procesessively by the ribosome-associated Rel. Rel dissociates from the ribosome either as a complex with tRNA or by itself after departure of deacylated tRNA. In the case of Rel:tRNA dissociating as a binary complex, the enzyme is expected to depart from the ribosome in a catalytically inactive HD^OFF^ SYNTH^OFF^ state.

**Figure 6.**
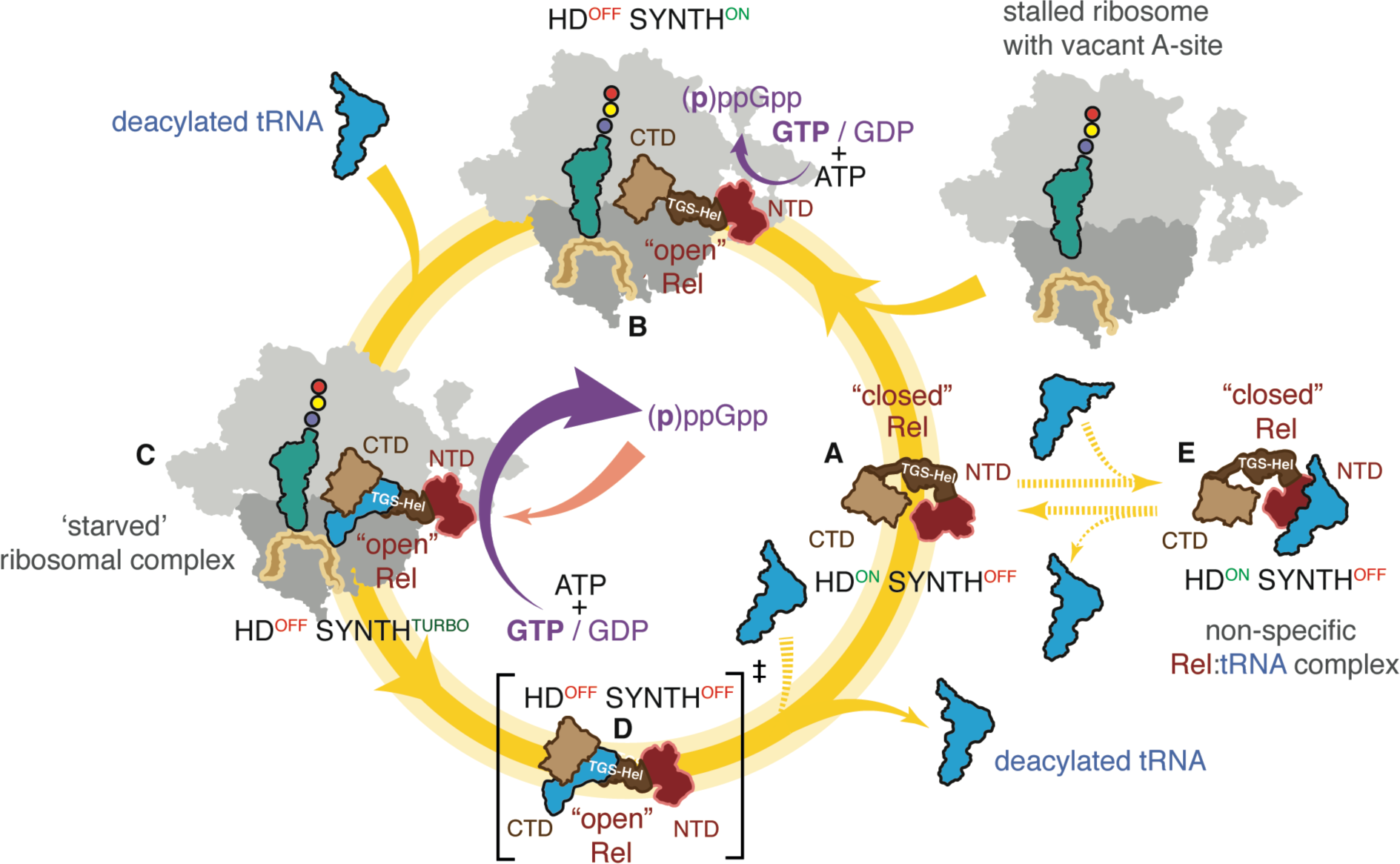
Model of Rel regulation by ‘starved’ ribosomal complexes. (**A**) Off the ribosome Rel assumes a ‘closed’ conformation concealing the tRNA-binding TGS and Helical domains. The factor is in hydrolytically active (HD^ON^ SYNTH^OFF^). In the ‘closed’ conformation the NTD region non-specifically binds tRNA (**E**). Upon binding to the vacant A-site of a starved ribosome, Rel switches to an ‘open’ conformation which stimulates the (p)ppGpp synthesis and inhibits (p)ppGpp hydrolysis (HD^OFF^ SYNTH^ON^) (**B**). Opening up primes the factor for specific recognition of deacylated tRNA by Rel’s TGS and Helical domains. Binding of the deacylated tRNA stabilises Rel on the ribosome and leads to full activation of the processive (p)ppGpp synthesis activity (HD^OFF^ SYNTH^TURBO^) (**C**). When the RelA:tRNA complex falls off the ribosome, the protein is in a catalytically inactive state due to the tRNA inhibiting the hydrolysis activity of the ‘open’ Rel (**E**). Dashed lines signify interactions that are observed in biochemical assays but their physiological relevance is unclear.

Our study further nuances the understanding of how the enzymatic activities of the NTD region are regulated by the CTD region. It is well-established that the regulatory CTD region of the protein transduces the signals from the ribosome and the tRNA, whereas the catalytic NTD ‘core’ of the protein is controlled via internal antagonistic allosteric control connecting the SYNTH and HD domains (16,21). While the two enzymatic activities of the NTD are antagonistic, the inactive state of one of the domains does not automatically necessitate activation of the other. For instance, the inhibition of hydrolysis by tRNA does not induce the activity of the SYNTH domain. At the same time, induction of one enzymatic domain efficiently represses the other, thus preventing the non-productive idle cycles of synthesis and destruction of the alarmone. The main manifestation of this mechanism is suppression of the hydrolysis activity upon association with a vacant ribosomal A-site or a starved ribosomal complex. Finally, our comparative biochemical investigation demonstrates that while the CTD-mediated interaction with the starved ribosome is essential for full activation of the Rel SYNTH domain, the connection between the activity and ribosomal binding is somewhat ‘loose’, with a broad range of acceptable ribosomal affinities for long RSHs. While *E. coli* RelA is a weak tRNA binder with a correspondingly low affinity to starved ribosomes, *B. subtilis* Rel has a significantly higher tRNA affinity and is stably recruited to the ribosome upon amino acid starvation. We hypothesise that the stable high-affinity interaction of Rel with starved ribosomal complexes is essential for efficient suppression of the hydrolysis activity upon amino acid starvation. In the case of the synthesis-only RelA this fail-safe mechanism is not needed, and the protein is evolutionary optimised for efficient sampling of the ribosomal population though dissociation and re-binding.

## SUPPLEMENTARY DATA

*Supplementary Data* are available at bioRxiv online.

## Author contributions

VH, HT and AGP drafted the manuscript, HT, AGP and VH coordinated the study, HT, VH and AGP designed experiments and analysed the data, HT, MR, JC-M, SJ, PK, KVN, RM, KA and TF performed experiments.

## ACKNOWLEDGMENTS

We are grateful to Protein Expertise Platform (PEP) at Umeå University and Mikael Lindberg for constructing plasmids, Gemma C. Atkinson for insightful comments on the manuscript, Kenn Gerdes and Kristoffer Skovbo Winther for sharing MG1655 *relA::HTF E. coli* strain (25).

