## Supplementary Data for "Ribosome association primes the stringent factor Rel for recruitment of deacylated tRNA to ribosomal A-site"

for

### Supplementary methods

*Plasmid construction*

pET24d -based expression constructs for expression and purification of mutant variants of *B. subtilis* Rel and *E. coli* RelA fused with an N-terminal $His_{10}$-SUMO tag were constructed by the Protein Expertise Platform at Umeå University. Expression constructs for wild type *E. coli* RelA (1) and *B. subtilis* Rel (2) were described earlier.

*Preparation of polyclonal anti-Rel antiserum*

Preparation of anti-Rel antiserum was described earlier (2).

*Preparation of 10X Polymix buffer base*

The 10X Polymix base was prepared as per (2).

*Preparation of $^{3}$H-labelled ppGpp and pppGpp*

Preparation of $^{3}$H-labelled pppGpp was described earlier (2). $^{3}$H- ppGpp was prepared similarly, with minor modifications: 2 µM *E. fecalis* RelQ (3) was incubated in reaction buffer (8 mM $MgCl_2$, 20 mM DTT, 20 M Tris-HCl pH 8.0) together with 2 mM ATP and 0.5 mM $^{3}$H-GDP (PerkinElmer, specific activity ≈80 cpm/pmol) for 30 min at 37 °C to produce $^{3}$H-ppGpp. Subsequent purification steps were identical.

*Protein expression and purification*

B. subtilis *Rel used for enzymatic assays, EMSA and sucrose gradient centrifugation experiments*: Expression and puficiation was perforemd as described earlier (2). Gel filtration profiles for mutant Rel variants are shown as (**Supplementary Figure S1**).

B. subtilis *$Rel^{NTD}$ used for isothermal titration calorimetry measurements*: the protein was purified by the same method as mentioned above, and digested with 35 µg of $His_6$-Ulp1 per 1 mg of Rel as well as 200 times diluted RNAse A (Sigma Aldrich) overnight at 10 °C. To get rid of the cleaved $His_{10}$-SUMO tag the protein was then loaded on a HiLoad 16/600 Superdex 200 pg column pre-equilibrated with ITC buffer (25 mM HEPES pH 8, 720 mM KCl, 50 mM L-glutamate, 50 mM L-arginine, 2 mM $MgCl_2$, 2 mM $MnCl_2$, 2% glycerol). The fractions enriched $Rel^{NTD}$ were pooled and the sample was dialyzed in Slide-A-Lyser 10 000 MWCO Dialysis Cassettes (ThermoScientific) against the ITC buffer overnight at 4 °C. For ITC titrations of GDP and GTP into $Rel^{NTD}$ at various temperatures the protein was purified as described above but the ITC buffer did not contain 2 mM $MnCl_2$. Protein concentration was estimated using the theoretical extinction coefficient at 280 nm (ProtParam online tool) and the purity was assessed by SDS-PAGE and spectrophotometrically ($OD_{260}/OD_{280}$ ratio below 0.6).

E. coli *RelA used for biochemistry, wild type and mutant variants*: all proteins were expressed, purified and characterised as described earlier (1).

*Preparation of 70S ribosomes*

*B. subtilis* 70S were purified as rescribed earlier (2). *E. coli* 70S ribosomes were prepared from the MRE600 strain (4) as described for *B. subtilis* 70S.

*Preparation of 70S initiation complexes (70S IC)*

*B. subtilis* 70S IC were purified as rescribed earlier (2).

*tRNA aminoacylation and formylation*

*E. coli* fMet-$tRNA_i^{Met}$ used to prepare *B. subtilis* 70S initiation complexes was prepared and purified as previously described (5) with minor modifications. Speficially, $^{3}$H-Methionine from PerkinElmer was use instead of $^{35}$S-Methionine._Aminoacylated $tRNA_i^{Met}$ and $tRNA^{Val}$ used for EMSA assays were prepared as previously described for fMet-$tRNA_i^{Met}$ (5) with N10-formyltetrahydrofolate and methionyl-tRNA formyltransferase (FMT) omitted from the reaction mix. Methionyl-tRNA synthetase and Valyl-tRNA synthetase were used for acylation of $tRNA_i^{Met}$ and $tRNA^{Val}$, respectively.

*Biochemical assays*

Synthesis assays were performed as described earlier for *E. coli* RelA and *B. subtilis* Rel (2), using either 300 μM $^{3}$H-GTP or 300 μM $^{3}$H-GDP (both from PerkinElmer) as well as 1 mM ATP as substrates. Hydrolysis *assays* were performed using either 300 μM $^{3}$H-pppGpp or 300 μM $^{3}$H-ppGpp as substate as rescribed earlier (2).

*Sucrose gradient fractionation and Western blotting*

*Sample preparation i: lysates*: *B. subtilis* samples were prepared as rescribed earlier (2) and supplemented with nucleotides GTP / GDP (0.5 mM) and ATP / AMPCPP (1 mM) as indicated on the figure legends and used for analytical ultracentrifugation and Western blotting.

*Sample preparation ii: lysates*: MG1655 *relA::HTF E. coli* strain (6) was grown in liquid LB media at 37 °C. At $OD_{600}$ of 0.2 the culture was treated 20 min with mupirocin added to final concentration of 70 μM, and the samples were processed as described for *B. subtilis* lysates.

*Sample preparation iii: reconstituted system*: 20 μL reaction mixtures containing 500 nM *B. subtilis* 70S IC(MVF), 140 nM Rel, *E. coli* $tRNA^{Val}$ (all in HEPES:Polymix buffer, 5 mM $Mg^{2+}$ final concentration) were incubated at 37 °C for 5 minutes and loaded onto sucrose gradients.

*Sucrose gradient fractionation*: clarified cell lysates (or reconstituted ribosomal complexes) were loaded onto 10–35% sucrose gradients in HEPES:Polymix buffer pH 7.5 (5 mM $Mg^{2+}$ final concentration, supplemented with nucleotides GTP:$Mg^{2+}$ / GDP:$Mg^{2+}$ (0.5 mM) and ATP:$Mg^{2+}$ / AMPCPP:$Mg^{2+}$ (1 mM) as indicated on the figure legends), subjected to centrifugation (36,000 rpm for 3 hours at 4 °C, SW-41Ti Beckman Coulter rotor) and analysed using Biocomp Gradient Station (BioComp Instruments) with $A_{260}$ and $A_{280}$ as a readout.

*Western blotting*: experiemtns were perforemd as described earlier (2). Ribosomal protein L3 of the 50S ribosomal subunit was detected using anti-L3 primary antibodies (a gift from Fujio Kawamura (7)) combined with goat anti-rabbit IgG-HRP secondary antibodies.

*Electrophoretic Mobility Shift Assay (EMSA)*

Before performing the experiment stock mRNA(MVF) (5'-GGC**AAGGAGGA**GAUAAGAAUGGUUUUCUAAUA-3') was incubated for 2 min at 60 °C to denature possible secondary structures. Reaction mixtures (10 µL) in HEPES:Polymix buffer (8) with 5 mM $Mg^{2+}$ were assembled by adding *E. coli* either $tRNA^{Val}$ / $tRNA_i^{Met}$ (0.1 µM final concentration) or mRNA (0.1 µM final concentration) or both (0.1 µM $tRNA^{Val}$ / $tRNA_i^{Met}$ and 1 µM mRNA(MVF) competitor), followed by the addition of Rel. After incubation for 5 min at 37 °C, 4 µL of 50% sucrose was added per sample, and the samples were electrophoretically resolved on a 12% Tris:Borate:EDTA gel at 4 °C (160-180V) for 1-1.5 h. Gels were stained with SYBR Gold nucleic acid stain (Life Technologies) for 30 minutes, followed by visualization using a Typhoon Trio Variable Mode Imager (Amersham Biosciences). Bands were quantified using ImageQuant TL (Amersham Biosciences). The efficiency of complex formation (effective concentration, $EC_{50}$) was calculated using the 4PL model (Hill equation) as per Sebaugh (9).

*Isothermal Titration Calorimetry (ITC)*

Dialyzed *B. subtilis* $Rel^{NTD}$ / *E. coli* $RelA^{NTD}$ was concentrated in Sartorius Ultrafiltration Centrifugal Concentrators (cut-off 30 kDa) at 3000 x g to a final concentration of 35 µM. 10 mM ppGpp and pppGpp (both from Jena Bioscience) were diluted to 450 µM in ITC buffer. Non-hydrolysable ATP analogue AMPCPP (ApCpp, Jena BioScience) and GDP were diluted with ITC buffer to 100 µM and 100 µM, respectively, and mixed with the protein prior to the experiment. 300 µM EDTA was incubated with *B. subtilis* $Rel^{NTD}$ overnight at 4 °C and supplemented with AMPCPP and GDP prior to the experiment. The samples were degassed and equilibrated at titration temperature and the ITC measurement were performed using Affinity ITC calorimeter (TA instruments) at 20 °C. The stirring rate was set to 75 rpm and a constant injection volume of 2 µL of titrant (ppGpp or pppGpp) was injected into the cell (177 µL of *B. subtilis* $Rel^{NTD}$) with an injection interval time of 250 seconds. For the heat exchange measurements $Rel^{NTD}$ protein was concentrated to 150 µM and 2.1 mM stock solutions of GDP and GTP were used. The stirring rate was set to 300 rpm and the measurements were performed at 20, 25 and 30 °C. All data were processed and analysed using the NanoAnalyse and Origin software packages. **Supplementary Figure S11** shows the representative results for each titration.

**Supplementary Table 1. Strains used in this work.**

| Strain | Description | Reference |
|---|---|---|
| *E. coli* | | |
| BL21 DE | B F⁻ *ompT gal dcm lon hsdS*$_B$(*r*$_B^-$*m*$_B^-$) λ(DE3 [*lacI lacUV5-T7p07 ind1 sam7 nin5*]) [*malB*$^+$]$_{K-12}$(λ$^S$) | Laboratory stock |
| MG1655 *relA::HTF* | *relA::HTF* | (6) |
| *B. subtilis* | | |
| wild type 168 | *trpC2* | Laboratory stock |
| VHB47 | *trpC2 rplK::cmR* | (2) |
| VHB68 | *trpC2 relH420E-spcR* | (2) |
| VHB156 | *trpC2 amyE::Phy-spnak-relD264G cmR rel::ermR* | (2) |
| RIK2508 | *trpC2* Δ*hpf* | (10) |

**Supplementary Table 2. Plasmids used in this work.**

| Plasmid | Description | Reference |
|---|---|---|
| VHP186 | pET24d-*$His_{10}$-SUMO-rel kmR* | (2) |
| VHP187 | pET24d-*$His_{10}$-SUMO-relD264G kmR* | this work |
| VHP230 | pET24d-*$His_{10}$-SUMO-relH420E kmR* | this work |
| VHP231 | pET24d-*$His_{10}$-SUMO-$rel^{\Delta RRM}$ kmR* | (2) |
| VHP232 | pET24d-*$His_{10}$-SUMO-$rel^{\Delta ZFD\text{-}RRM}$ kmR* | this work |
| VHP233 | pET24d-*$His_{10}$-SUMO-$rel(1\text{-}373)^{NTD}$ kmR* | this work |
| VHP596 | pET24d-*$His_{10}$-SUMO-$rel^{TGS\text{-}Helical}$ kmR* | this work |
| VHP597 | pET24d-*$His_{10}$-SUMO-$rel^{TGS}$ kmR* | this work |
| VHP598 | pET24d-*$His_{10}$-SUMO-$rel^{TGSH420E\text{-}Helical}$ kmR* | this work |
| VHP599 | pET24d-*$His_{10}$-SUMO-$rel^{TGSH420E}$ kmR* | this work |

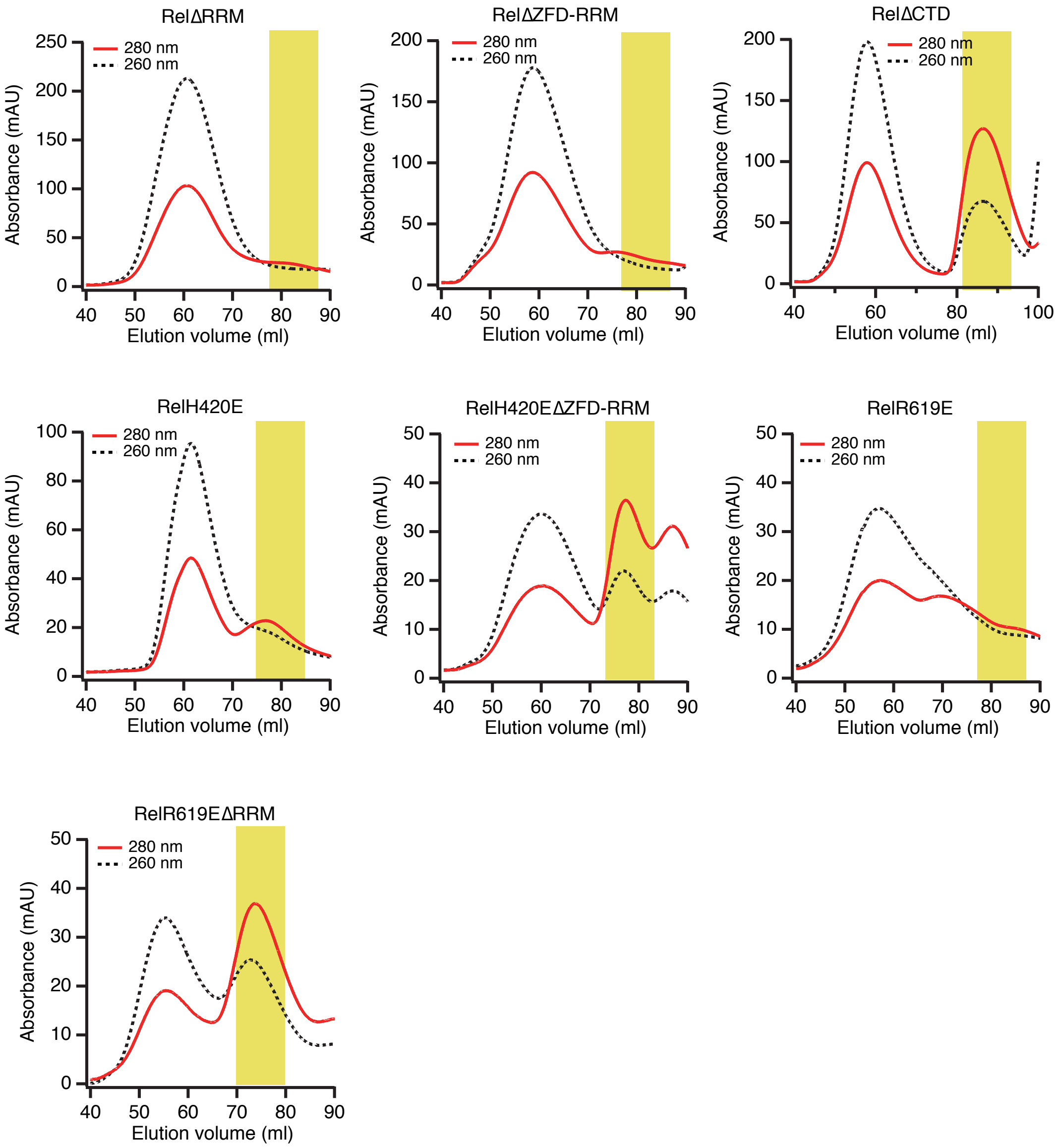


**Supplementary Figure S1. Gel filtration purification step of *B. subtilis* Rel mutants using HiLoad 16/600 Superdex 200 pg.** Only fractions highlighted in yellow were carried out to the next purification step. Gel filtration profile of ΔRRM Rel is adapted from (2).

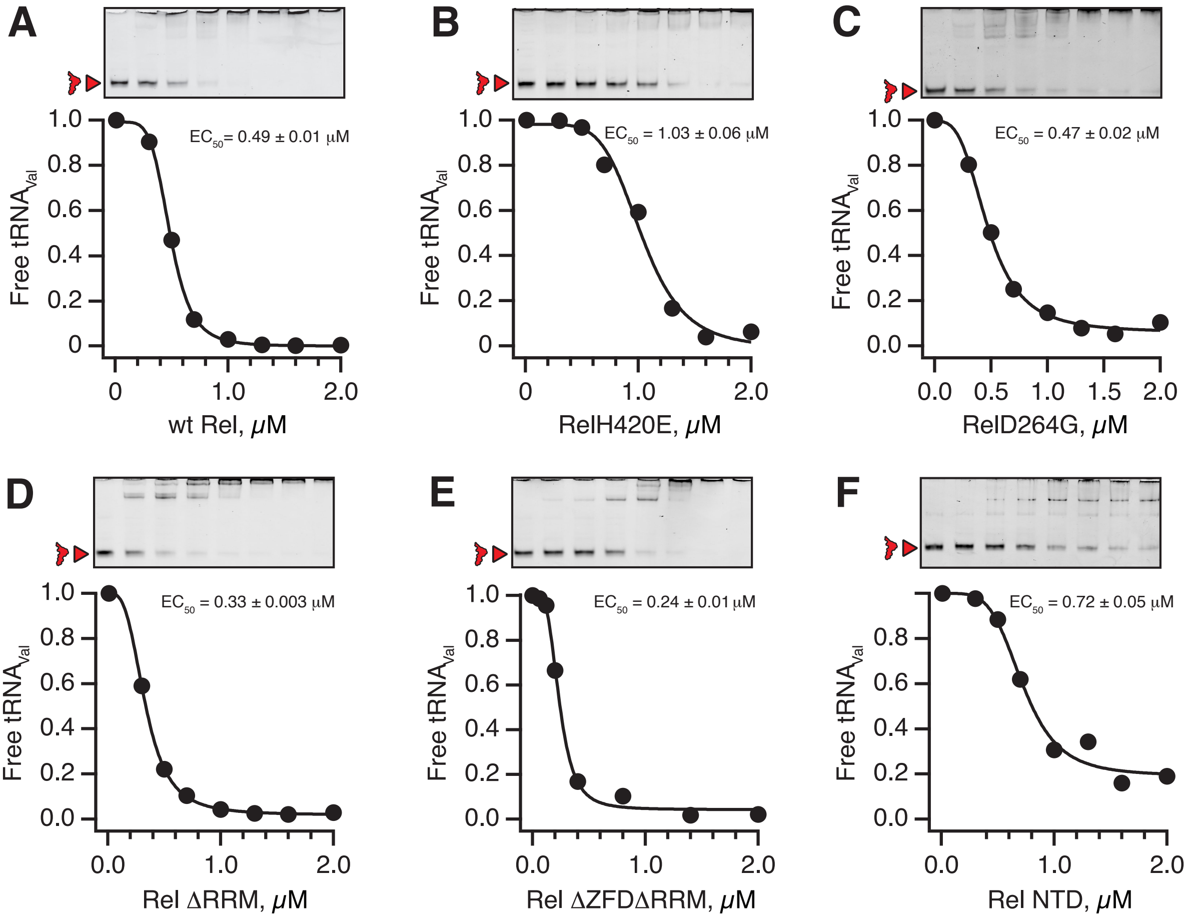


**Supplementary Figure S2. *B. subtilis* Rel complex formation with *E. coli* tRNA$^{Val}$ in the absence of non-specifric RNA competitor.** Complex formation between 0.1 µM *E. coli* tRNA$^{Val}$ and increasing concentrations of *B. subtilis* Rel (wt or mutants) was monitored by EMSA in the absence of competing mRNA(MVF). The efficiency of complex formation (effective concentration, $EC_{50}$) was calculated using the 4PL model (Hill equation) as per Sebaugh (9).

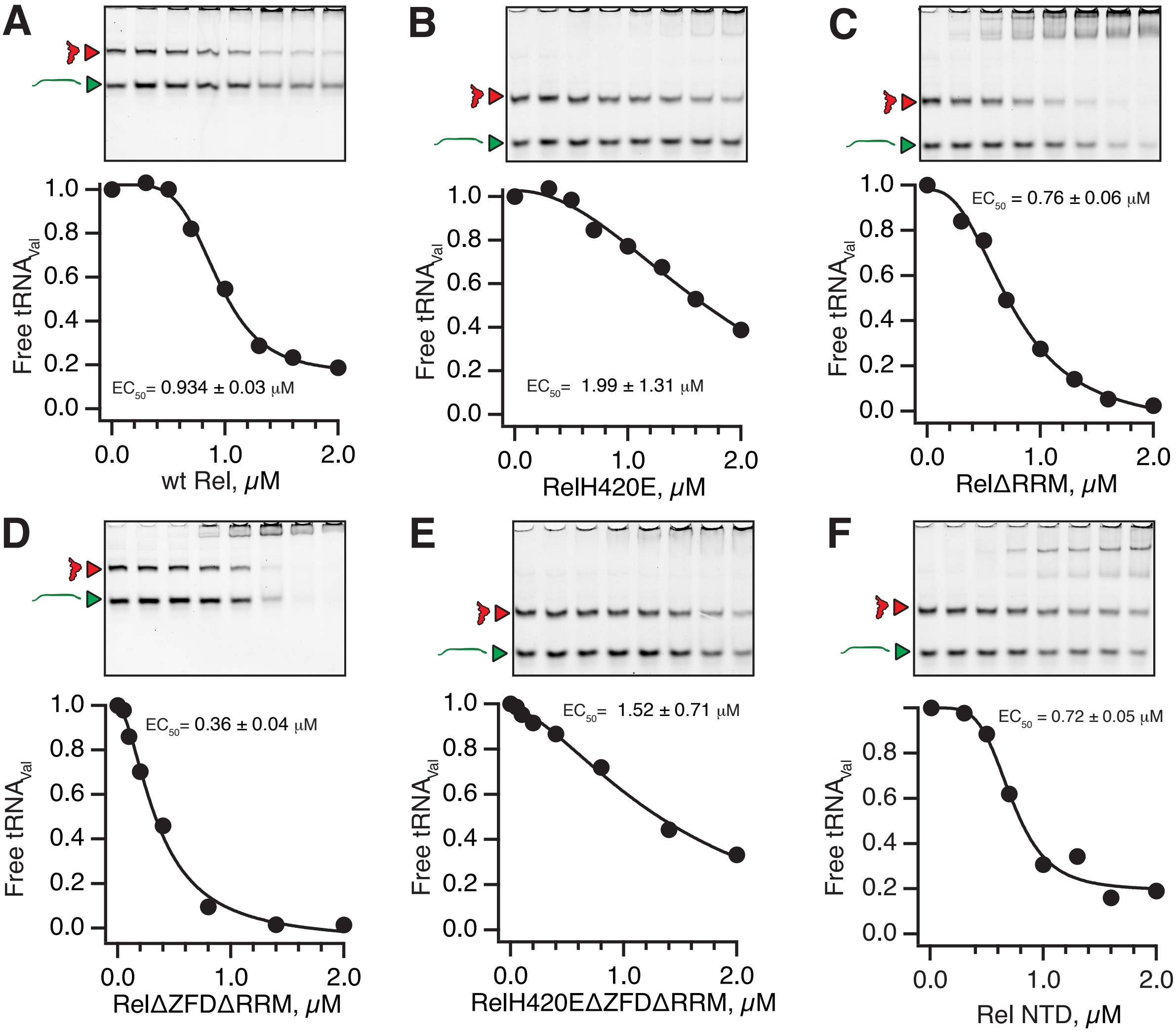


**Supplementary Figure S3. *B. subtilis* Rel complex formation with *E. coli* tRNA$^{Val}$ in the presence of non-specifric RNA competitor.** Complex formation between 0.1 µM *E. coli* tRNA$^{Val}$ and increasing concentrations of *B. subtilis* Rel (wt or mutants) was monitored by EMSA in the presence of competing mRNA(MVF) (1 µM). The efficiency of complex formation (effective concentration, $EC_{50}$) was calculated using the 4PL model (Hill equation) as per Sebaugh (9).

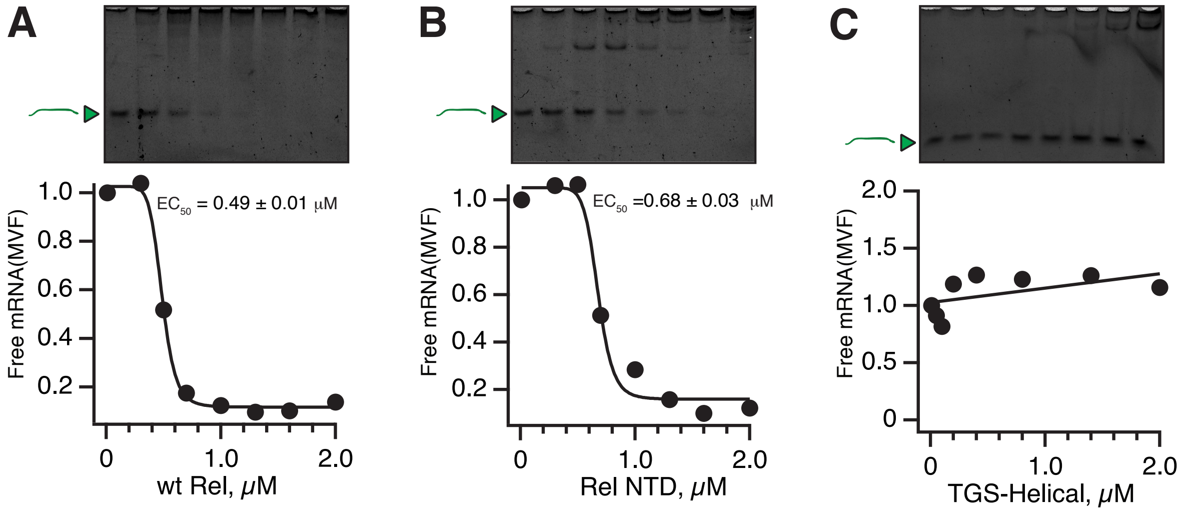


**Supplementary Figure S4.** ***B. subtilis*** **Rel NTD mediates non-specific RNA binding.** Complex formation between 0.1 μM mRNA(MVF) and increasing concentrations of *B. subtilis* Rel (wt or mutants) was monitored by EMSA. The efficiency of complex formation (effective concentration, $EC_{50}$) was calculated using the 4PL model (Hill equation) as per Sebaugh (9).

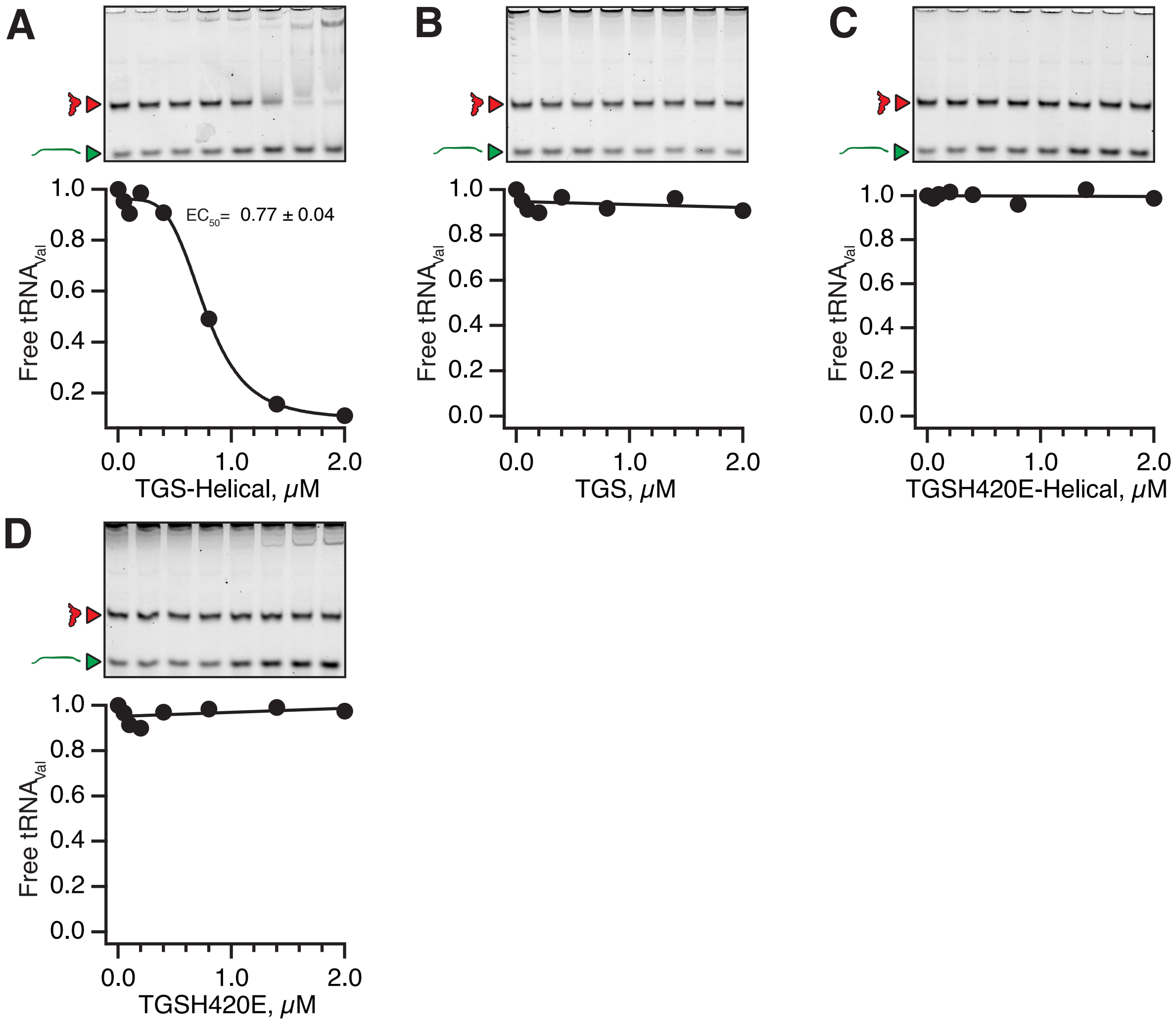


**Supplementary Figure S5. Specific complex formation between *B. subtilis* Rel TGS-Helical fragment and *E. coli* tRNA$^{Val}$.** Complex formation between 0.1 μM *E. coli* tRNA$^{Val}$ and increasing concentrations of *B. subtilis* Rel fragments was monitored by EMSA in the presence of competing mRNA(MVF) (1 μM). The efficiency of complex formation (effective concentration, $EC_{50}$) was calculated using the 4PL model (Hill equation) as per Sebaugh (9).

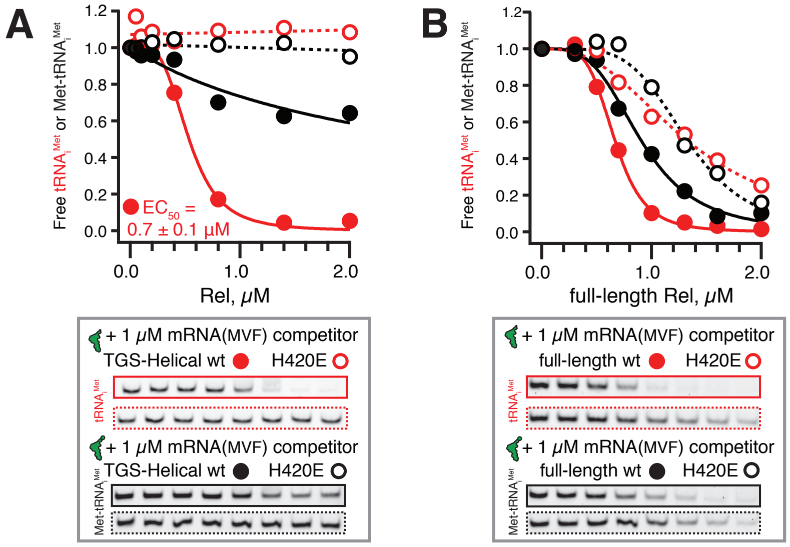


**Supplementary Figure S6. Complex formation between *B. subtilis* Rel TGS-Helical fragment and *E. coli* tRNA$_i^{fMet}$ is abrogated by tRNA$_i^{Met}$ aminoacylation.** Complex formation between 0.1 µM initiator tRNA$_i^{fMet}$ and increasing concentrations of *B. subtilis* Rel (either TGS-Helical region (**A**) or full-length (**B**), wt (filled cyrcles) or H420E mutants (empty circles)) was monitored by EMSA. The efficiency of complex formation (effective concentration, $EC_{50}$) was calculated using the 4PL model (Hill equation) as per Sebaugh (9).

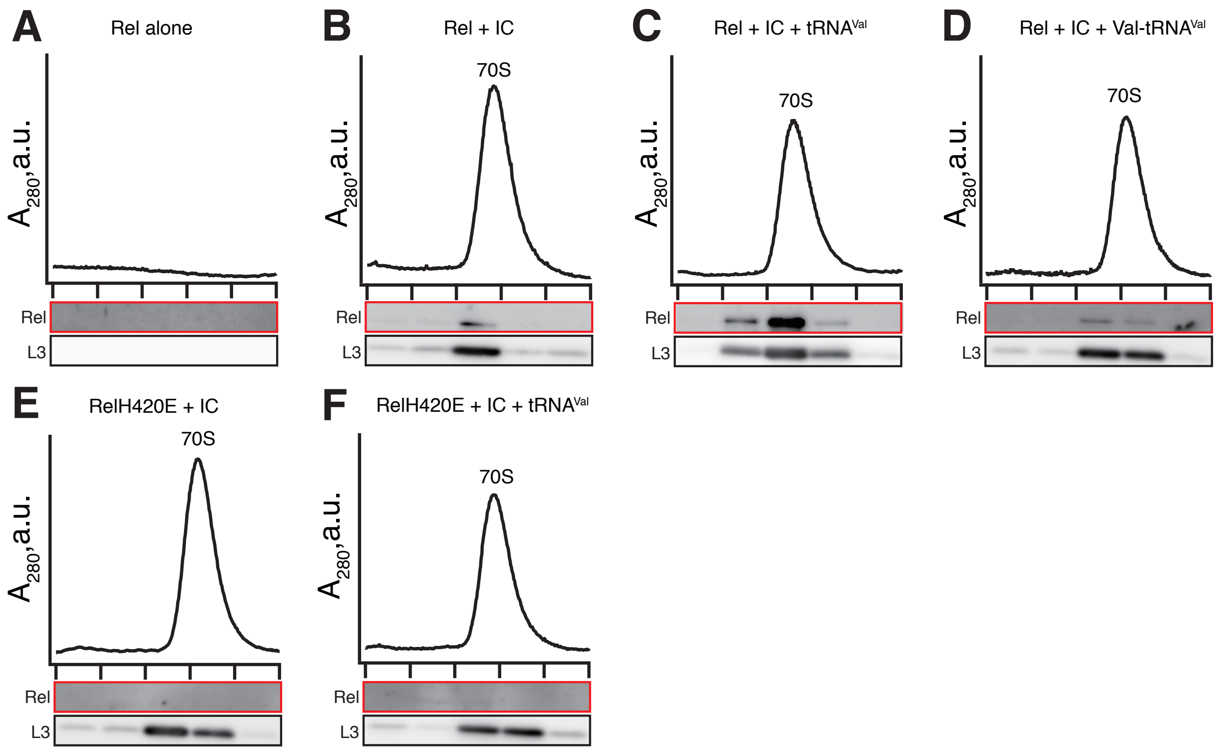


**Supplementary Figure S7. Sucrose gradient centrifugation and immunobloting analyses of reconstituted Rel:ribosome complexes.** As indicated on the figure, 25 µl of 140 nM either Rel protein (**A**) or RelH420E protein mixtures were supplemented with combinations of 0.5 µM IC (MV) (**B** and **E**) and either tRNA$^{Val}$ (**C** and **F**) or Val-tRNA$^{Val}$ (2 µM: A-site) (**D**) were incubated at 37 °C for 5 min and resolved on 10 to 35% sucrose gradients in HEPES:Polymix buffer (pH 7.5, 5 mM $Mg^{2+}$). All experiments were performed in HEPES:Polymix buffer, pH 7.5 at 37 ˚C in the presence of 5 mM $Mg^{2+}$.

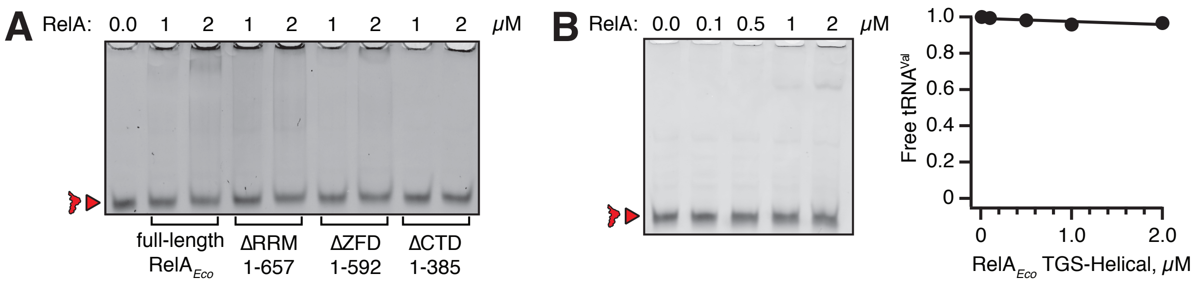


**Supplementary Figure S8. No complex formation is detected by EMSA between *E. coli* Rel TGS-Helical fragment and *E. coli* tRNA$^{Val}$.** Complex formation between 0.1 μM *E. coli* tRNA$^{Val}$ and *E. coli* RelA, full-length or C-terminally truncated (**A**) as well as isolated *E. coli* RelA TGS-Helical region (**B**) was monitored by EMSA.

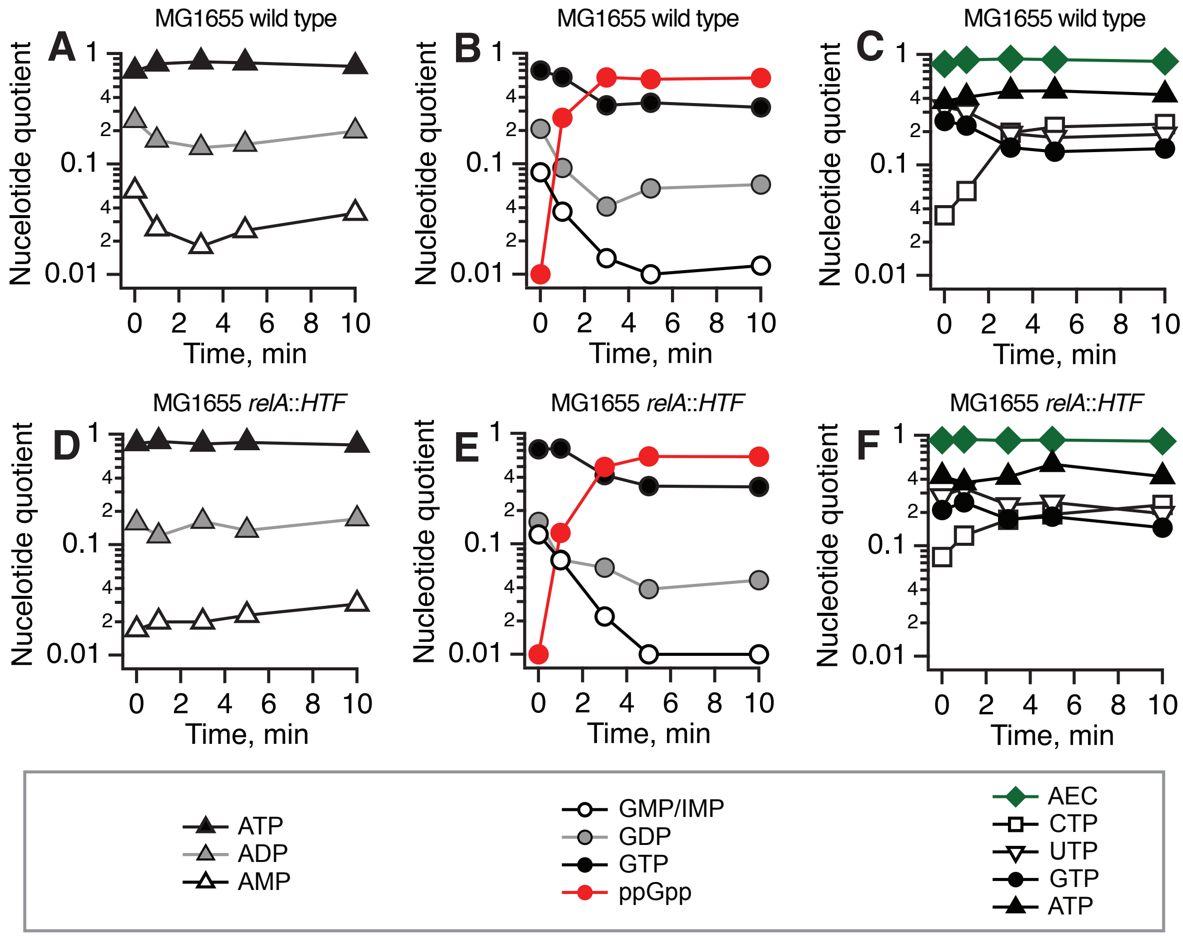


**Supplementary Figure S9. C-terminally HTF-tagged *E. coli* RelA displays wild type-like functionality in live *E. coli*.** Wild type MG1655 and MG1655 *relA::HTF* (6) *E. coli* strains were grown in MOPS minimal media at 37 °C until $OD_{600}$ 0.5 and challenged with 150 µg/mL of mupirocin (3X the MIC). HPLC-based nucleotide quantification was performed as per Varik and colleagues (11).

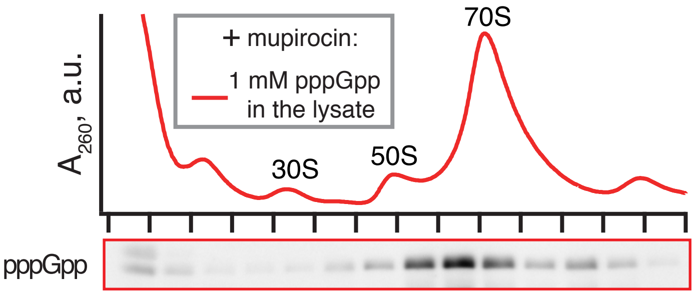


**Supplementary Figure S10. Addition of pppGpp to lysate does not destabilise Rel's association with starved ribosomes.** Wild type Rel was expressed in wild type 168 *B. subtilis.* To induce amino acid starvation exponentially growing bacterial cultures at $OD_{600}$ 0.2 were treated with mupirocin for 20 min. The drug was added to a final concentration of 700 nM that completely abolished the growth. The lysates were supplemented with 1 mM pppGpp (final concentration) and then resolved on 10 to 35% sucrose gradients in HEPES:Polymix buffer (pH 7.5, 5 mM $Mg^{2+}$) supplemented with nucleotides as indicated on the figure (GTP:$Mg^{2+}$ and GDP:$Mg^{2+}$ at 0.5 mM; ATP:$Mg^{2+}$ and AMPCPP:$Mg^{2+}$ at 1 mM).

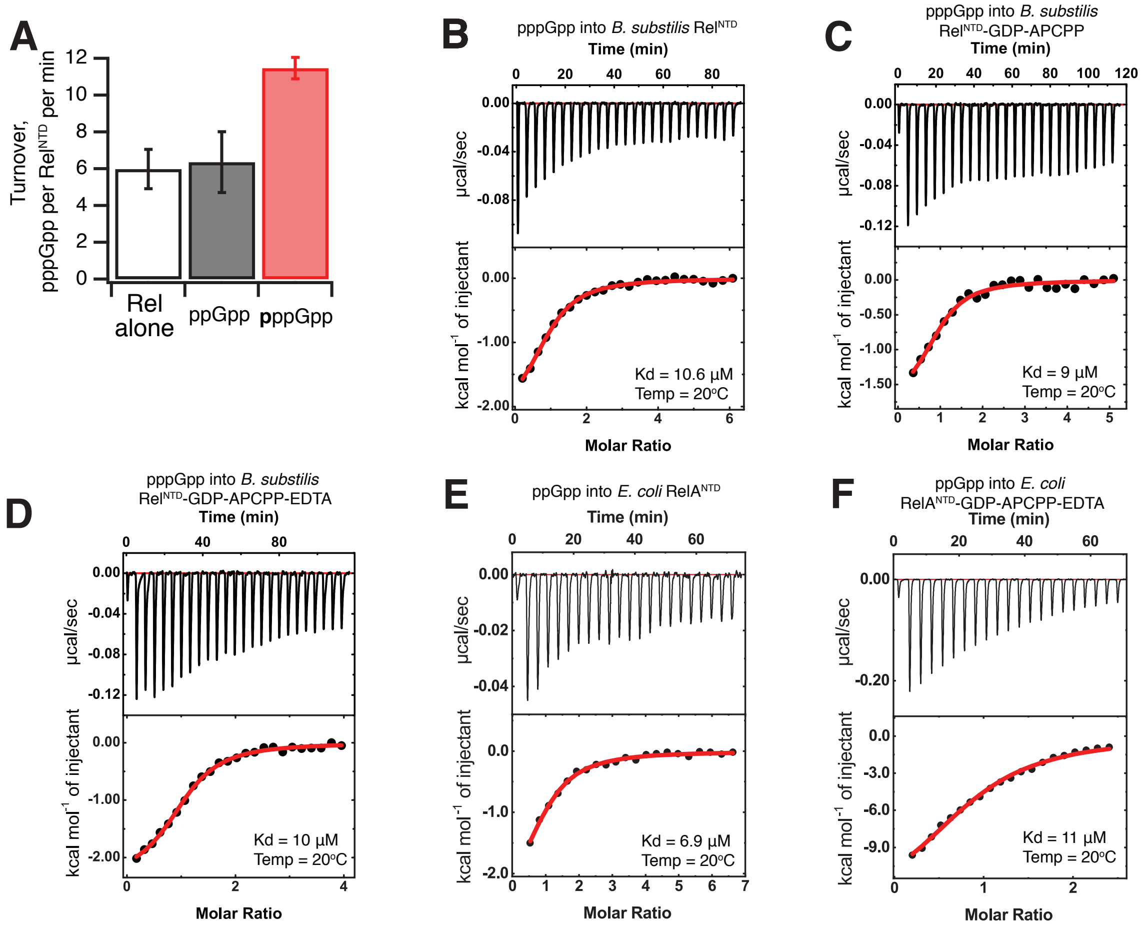


**Supplementary Figure S11. NTD domain resion of *B. subtilis* Rel and *E. coli* RelA contains the allosteric regulatory site that binds pppGpp.** (**A**) Synthesis activity of *B. subtilis* Rel$^{NTD}$ is promoted by 100 µM pppGpp but not ppGpp. pppGpp binds to *B. subtilis* Rel$^{NTD}$ (**B**), and the interaction is insensitive to addition of GDP and non-hydrolysable ATP analogue APCPP that occupy the SYNTH active site (**C**), or preincubation of Rel$^{NTD}$ with EDTA which removes the $Mn^{2+}$ ion from the catalytic site of the HD domain in the combination of GDP with EDTA pre-treatment (**D**). pppGpp binds to *E. coli* RelA$^{NTD}$ (**E**), and the interaction is insensitive to addition of GDP and non-hydrolysable ATP analogue APCPP and preincubation of with EDTA (**F**). Experiments were performed using 35 µM Rel$^{NTD}$, 450 µM pppGpp, 100 µM APCPP, 100 µM GDP and 300 µM EDTA.

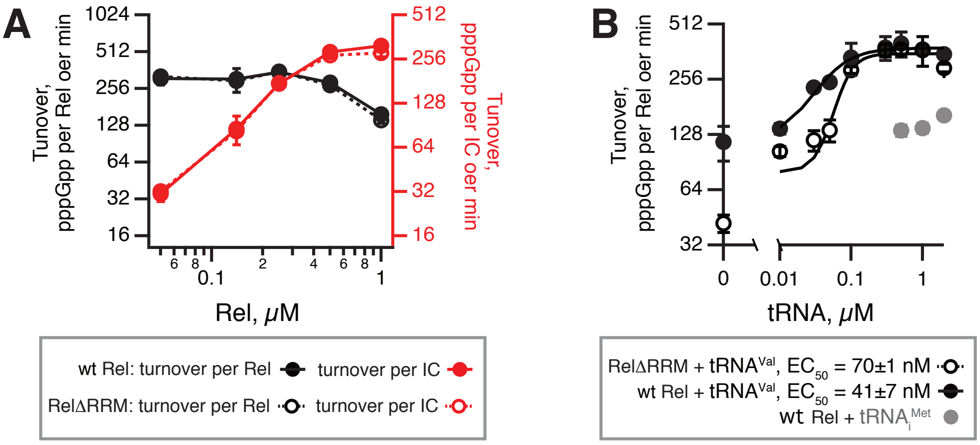


**Supplementary Figure S12. Synthase activity of full-length and Rel$^{\Delta RRM}$ as a function increasing concentrations of either Rel** (**A**) **or tRNA$^{Val}$** (**B**). All biochemical reaction mixtures contained 1 mM ATP, 0.3 mM $^{3}$H-labelled GTP and 0.5 μM 70S IC(MVF) as well as either increasing concentrations Rel in the presence of 2 μM tRNA$^{Val}$ (**A**) or increasing concentrations tRNA$^{Val}$ in the presence of 0.1 μM Rel (**B**). The error bars represent standard deviations of the turnover estimates by linear regression.
